# Transformations of cognitive maps for sensorimotor control

**DOI:** 10.64898/2026.03.25.714326

**Authors:** Joonhee Leo Lee, Yixuan Wang, Agostina Casamento-Moran, Kosisochukwu Ugorji, Jose Jarquin, Daniel C McNamee, Vikram S Chib

## Abstract

Adaptive embodied behavior involves transforming structured knowledge about the relationship between environment and action into motor signals, but how these transformations are coordinated across brain networks remain unknown. Participants learned associations between visual cues and isometric exertions that varied in force and duration, forming a two-dimensional cognitive map of a force–time space. During behavior, this force-time space was expressed in several cortical regions using grid-like coding schemes, indicating sensorimotor cognitive maps. Importantly, while mnemonic regions such as the entorhinal cortex maintained an unwarped, task-relevant representation, the primary motor cortex encoded a force-time space distorted by perceived effort during motor execution. Dynamic causal modeling showed inhibitory motor-to-mnemonic coupling that predicted the transformation of effort-weighted motor signals into sensorimotor maps. Furthermore, individual differences in learning and navigating the force-time space independently shaped mnemonic map geometry and perceived effort. These findings demonstrate that sensorimotor cognitive maps emerge from dynamic interactions between motor and mnemonic systems and are shaped by individual differences during the learning and execution of movement.

## Introduction

Skilled movement emerges from the adaptive coordination of motor actions with environmental information and sensory feedback. For example, learning to play the violin involves associating visual cues from sheet music with precise sensorimotor commands that determine the force and timing of bow movements. Motor commands are inherently embodied, shaped by high-dimensional and continuous parameters governing body physics such as timing^1^, position^2^, velocity^3^, and force^4,5^. Although population dynamics in sensorimotor regions such as primary motor cortex reflect such endogenous motor variables^1,4,6–9^, successful behavior also depends on representing actions in relation to exogenous sensory variables such as environmental cues in closed-loop^10^. While the nervous system readily forms cue–movement associations^11,12^, how sensorimotor information is encoded in the brain to relate motor parameters to exogenous environmental structure, and how this information is coordinated across brain networks to guide behavior remains unclear.

One potential framework for organizing such cue-movement information is cognitive maps. Cognitive map theory proposes that the brain constructs internal representations of structured relationships between components of an environment or ontology to support memory and guide adaptive behavior^13^. Such map-like coding has been observed within the brain’s mnemonic network, including the hippocampal formation^14–19^, which consists of the hippocampus and entorhinal cortex, and the retrosplenial cortex^20–25^. Although originally characterized in spatial navigation, cognitive map-like representations have also been reported in non-spatial domains^25^, such as social^26–28^ and conceptual spaces^29^, suggesting a more general role in organizing relational knowledge. These previous works have focused on how cognitive maps of exogenous features are constructed and organized by the brain. However, in the context of sensorimotor control, motor features are inherently endogenous to the body, originating from an individual’s embodied intentions and internal state. Whether such cognitive mapping principles apply to endogenous sensorimotor features remains unknown. If mnemonic systems construct sensorimotor cognitive maps that link movements to task-relevant goals, they must therefore receive and transform movement-related signals originating in sensorimotor regions, essentially forming relational knowledge between an individual’s endogenous sense of motor features and exogenous information about goals.

A central challenge for encoding sensorimotor cognitive maps is that endogenous movement representations may not faithfully reflect the task-relevant action space. For example, variations in force and timing of movements can differentially influence perceived effort during movement execution^6,30–32^. This motivates our hypothesis that perceived effort during movements can warp the representational geometry of endogenous movement features. If mnemonic systems construct task-relevant maps of action space, they must transform these effort-weighted endogenous motor signals into representations aligned with exogenous task demands. Whether, and how, interactions between mnemonic and sensorimotor networks support such transformations remain unknown.

Here, we investigated whether the human brain constructs a structured, cognitive map-like representation of a learned sensorimotor space and how this representation emerges through interactions between sensorimotor and mnemonic systems. Participants learned to associate visual cues with isometric hand-grip exertions that varied systematically in force and time (**Fig. 1A**). Using functional magnetic resonance imaging (fMRI), we tested whether the force and time of these cue-dependent exertions are represented within a force-time cognitive map across sensorimotor and mnemonic brain regions.

**Figure 1:**
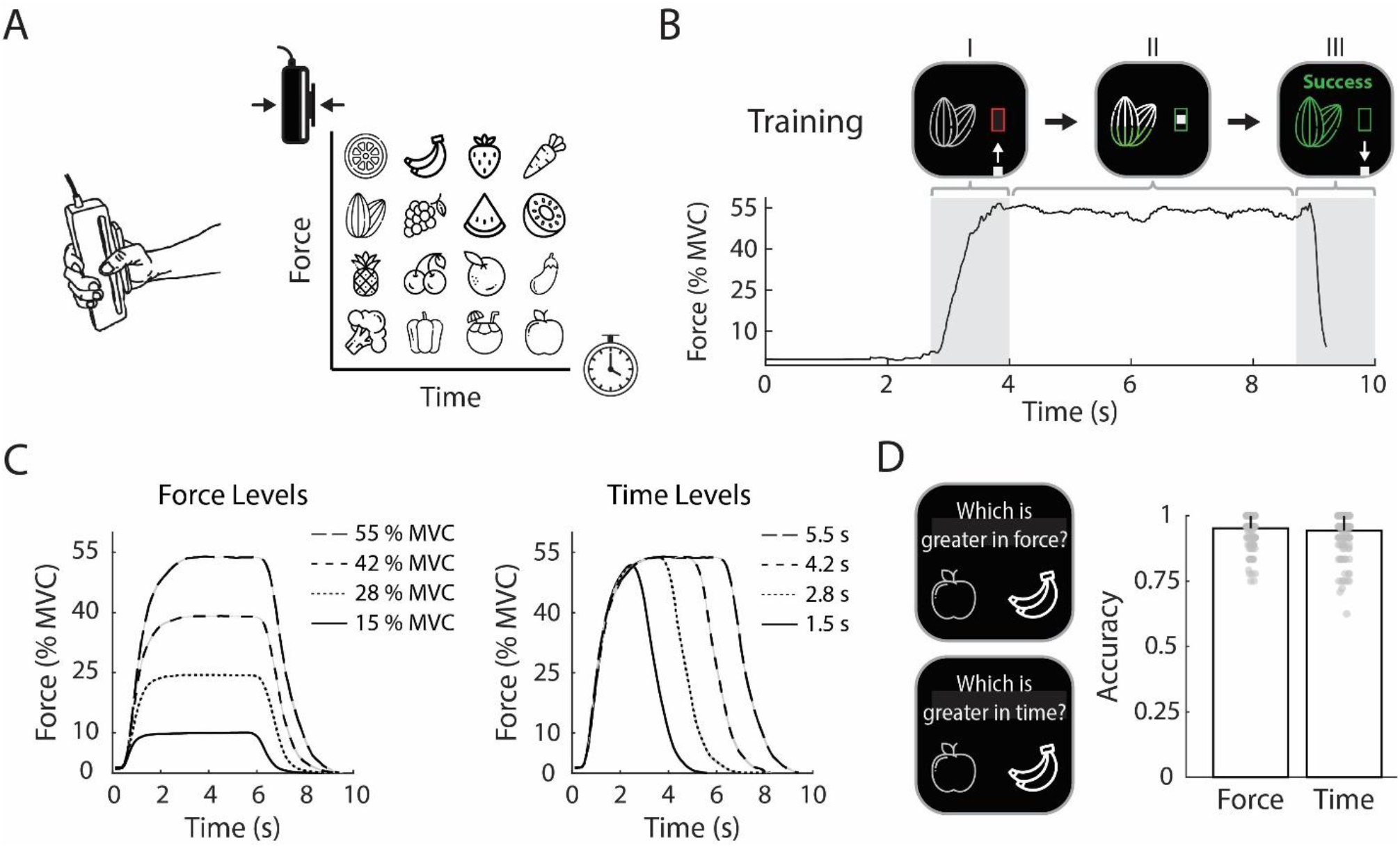
Participants associated effortful isometric exertions with external cues. **(A)** During training, participants performed isometric exertions using a hand dynamometer (left). They associated the force (intensity) and time (duration) of each exertion with a visual cue. The visual cues were arranged in a 4×4 grid-like force-time space (right), with four levels of force and time, respectively. **(B)** For every trial in the training phase, participants initially performed an isometric exertion to guide a cursor to the target box (Panel I). Upon reaching the target, the cue began to fill in green color and participants had to continue exerting at the target force level (Panel II). When the cue finished filling, participants let go of the hand dynamometer, and the cursor returned to baseline (Panel III). Force levels were assigned % of the participant’s maximum voluntary contraction (MVC). **(C)** Mean exertion profiles for different force (left, target time 5.5 s) and time (right, target force %55 MVC) levels. **(D)** During training, participants were periodically shown two cues and were asked to choose the one that was greater in force or time level. Participants chose the correct cue that had greater force or time levels with high accuracy. Error bars represent one standard deviation (std).

During navigation of the learned force–time space, the entorhinal cortex exhibited grid-like coding consistent with a cognitive map-like organization. In contrast, the primary motor cortex encoded a warped map representation reflecting perceived effort. Despite this distortion, mnemonic regions preserved an unwarped, task-relevant geometry corresponding to learned cue–exertion associations. Dynamic causal modeling revealed inhibitory motor-to-mnemonic coupling that predicted the transformation of effort-weighted signals into task-relevant maps. Moreover, individual differences in learning and effort perception independently shaped mnemonic map geometry. Together, these findings indicate that sensorimotor cognitive maps arise through dynamic interactions between sensorimotor and mnemonic systems, revealing how endogenous sensorimotor information is transformed into structured knowledge that supports goal-directed behavior.

## Results

### Participants associated effortful isometric exertions with external cues

Healthy human participants (n = 33) completed a multi-day study that assessed the cognitive map of structural knowledge in humans^26,33^. Participants learned to associate visual cues with effortful isometric hand-grip exertions and were later tested on their ability to combine learned cue-exertion mappings with sensorimotor feedback. Each of 16 cues mapped onto a unique combination of force and time, forming a structured 4×4 force-time space (**Fig. 1A**). Force determined the magnitude of exertion needed to reach a target, whereas time determined how long the exertion had to be maintained (**Fig. 1B,C**). Force was scaled to each participant’s maximum voluntary contraction (MVC) and ranged from 15 to 55 % of MVC, whereas time ranged from 1.5-5.5 seconds.

During the first two training days, participants learned to associate the cues with one dimension of exertion, either force or time, followed by the combination of both dimensions on Day 3 (**Supplementary Fig. 1-3**). Throughout training, participants were also assessed on their cue-exertion mappings. Participants were shown two cues and instructed to select the cue with either a greater force or time level, determined based on the training dimension (**Fig. 1D**; **Supplementary Fig. 3**). Participants demonstrated high accuracy in distinguishing cues based on both force (mean ± SD accuracy = 0.94 ± 0.08) and time (mean ± SD accuracy = 0.93 ± 0.12) levels, confirming their ability to associate cues with the learned force-time space.

### Participants learn to navigate sensorimotor force-time space

Participants also performed a separate Motor-Space Navigation (MSN) task that assessed their ability to utilize the force-time space that was learned during training with sensorimotor feedback and consisted of two segments: 1) localization and 2) navigation. During localization, participants performed an isometric exertion without being told the cue identity (visualized as a generic circle) and were instructed to think of the cue that matched the exerted force and time level in theforce-time space (**Fig. 2A**, Green). In the navigation segment, a target cue appeared (**Fig. 2A**, Blue), and participants were instructed to traverse the force-time space and judge the distance from the localized cue to the target cue. This process was repeated a second time with the same hidden cue during localization, but a different target cue in the navigation segment. Finally, participants selected the target cue that was further from the localized cue in the force-time space (D1 vs D2, **Fig. 2A**).

**Figure 2:**
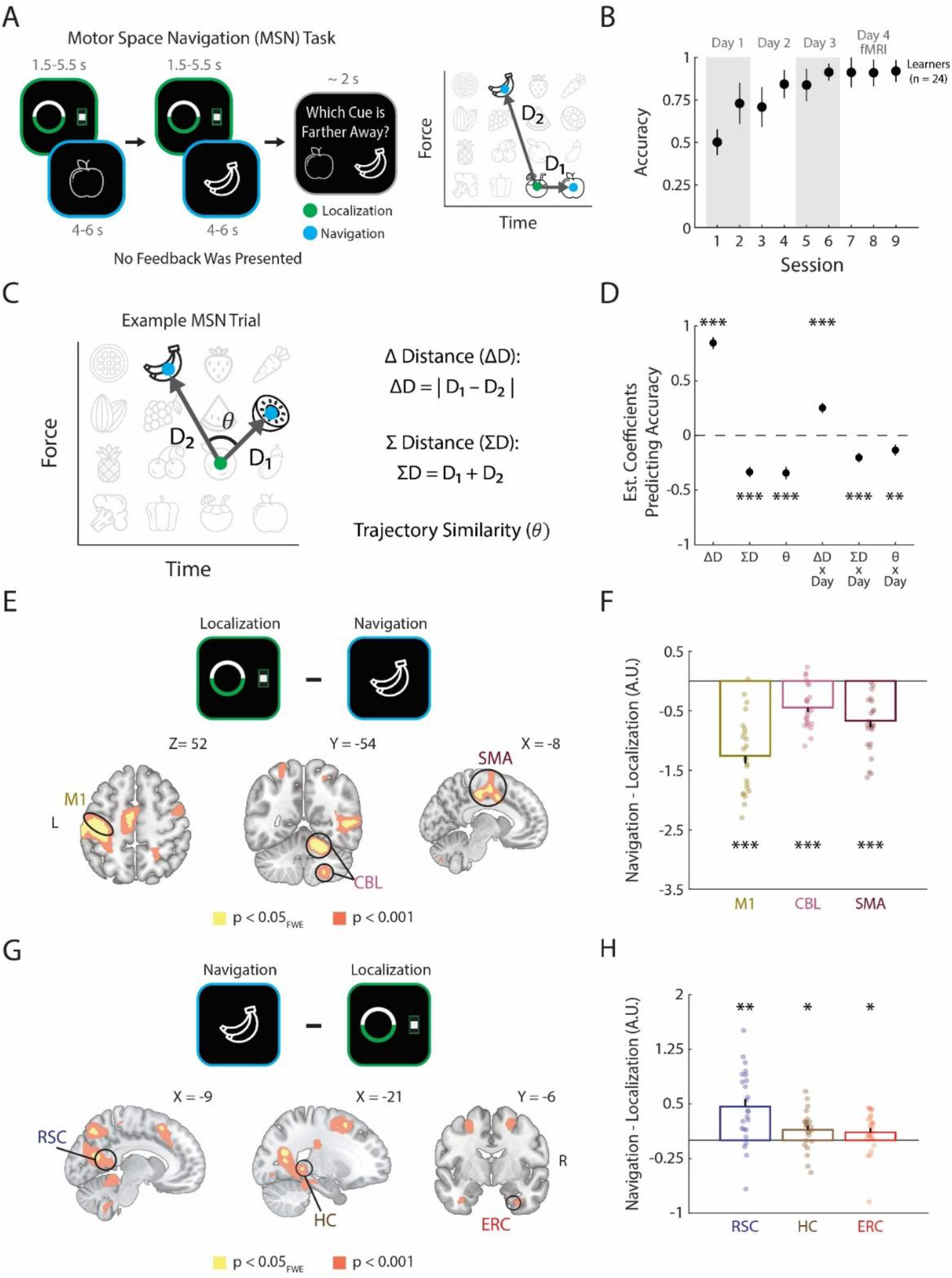
Participants learn to navigate sensorimotor force-time space. **(A)** During the motor space navigation (MSN) task, participants completed two segments: 1) localization and 2) navigation. During localization, participants performed an isometric exertion similar to what they did during training, but the cue was hidden and replaced with a circle. Participants had to identify the hidden cue that matched the exerted force and time level (green). During navigation, a target cue appeared, and participants had to compute the distances (D_1_, D_2_) between the hidden cue and the target cue in the force time space (blue). Finally, participants decided which target cue was farther from the hidden cue in the force-time space (D_1_ vs D_2_). **(B)** Participants’ mean performance in the MSN task across the four experimental days. Sessions 1-6 were throughout days 1-3. Participants who displayed greater than 80% accuracy on session 6 (n = 24) performed sessions 7-9 in the fMRI scanner. Error bars represent one std. **(C)** To investigate whether participants’ ability to navigate force-time space is spatially mediated, three metrics were computed from the trajectories that participants had to mentally traverse for each trial in the MSN task. Δ Distance (ΔD) was defined as the absolute difference between the Euclidean distance of two trajectories (|D_1_ – D_2_|). Σ Distance (Σ D) was defined as the sum of the Euclidean distance of two trajectories (D_1_ + D_2_). Trajectory similarity was defined as the angle (θ, [0, 180°)) between the two trajectories. **(D)** Estimated coefficients from a mixed-effects generalized linear model that predicts participants’ accuracy using the spatially mediated metrics, and interaction with experimental days. Error bars represent one standard error of the mean (sem). *** p < 0.001, ** p < 0.005 **(E)** Whole brain contrasts showing brain regions that were more active for localization than navigation (Localization - Navigation). Contrast is displayed at p < 0.05 FWE corrected (yellow) and p < 0.001 uncorrected (red). **(F)** Region of interest analysis (ROI) showed that the primary motor cortex (M1), cerebellum (CBL) and supplementary motor area (SMA) all showed significantly greater activity for localization than navigation segment of the MSN task. Error bars represent one sem. *** p < 0.001 two-tailed one-sample Wilcoxon signed-rank test, corrected for number of sensorimotor regions (n = 3) using Holm-Bonferroni method. **(G)** Whole brain contrasts showing brain regions that were more active for navigation than localization (Navigation - Localization). Contrast is displayed at p < 0.05 FWE corrected (yellow) and p < 0.001 uncorrected (red). **(H)** Region of interest analysis (ROI) showed that the retrosplenial cortex (RSC), hippocampus (HC) and entorhinal cortex (ERC) all showed significantly greater activity for navigation than localization segment of the MSN task. Error bars represent one sem. ** p < 0.005, * p < 0.05, two-tailed one-sample Wilcoxon signed-rank test, corrected for number of mnemonic regions (n = 3) using Holm-Bonferroni method.

Participants completed the MSN task before and after training across three behavioral testing days (**Supplementary Fig. 1**). Participants who passed our performance criteria (> 0.80 accuracy in MSN at the end of Day 3, session 6, n = 24) showed progressive improvement and performed the MSN task inside the fMRI scanner on the final day (**Fig. 2B**, **Supplementary Fig. 4D**). To further evaluate participants’ behavioral performance, participants performed a localization-only task, which tested the overt behavioral response following localization, and a spatial reconstruction task in which participants reconstructed the force-time space (SI **–** Additional Behavioral Assessments, **Supplementary Fig. 5-6**). Participants who met the performance criteria in the MSN task showed high accuracy in both tasks (**Supplementary Fig. 5-6)**, indicative of a stable representation of the learned force-time space.

To establish whether participant’s behavior reflected a cognitive map of force-time space, we modeled MSN accuracy using spatial metrics that measured differences in distance (ΔD), summed distance (ΣD), and trajectory similarity (θ) of the two trajectories for each trial (See **Methods**, **Fig. 2C**). Mixed-effects logistic regression revealed that ΔD was a significant positive predictor of MSN accuracy (β(SE) = 0.84(0.05), p < 0.001) (**Fig 2D, Supplementary Table 1**), indicating that participants were more likely to respond correctly when the difference between two trajectories was larger. In contrast, ΣD was a significant negative predictor of accuracy (β(SE) = -0.34(0.04), p < 0.001) (**Fig 2D**, **Supplementary Table 1**), suggesting that participants made more errors when they had to traverse greater distances in the force-time space. θ was a significant negative predictor (β(SE) = -0.34(0.05), p < 0.001) (**Fig 2D**, **Supplementary Table 1**), indicating reduced performance when the two trajectories diverged more in direction. All three spatial metrics exhibited significant interactions with experimental day (ΔD x Day β(SE) = 0.25(0.04); ΣD x Day β(SE) = -0.20(0.04); θ x Day β(SE) = -0.14(0.05); all p < 0.001) (**Fig 2D**, **Supplementary Table 1**), indicating that reliance on spatial structure strengthened with learning. Together, these results indicate that participants used a cognitive map-like representation of force-time space when performing the MSN task.

### Mnemonic and sensorimotor regions are engaged at distinct time points during the MSN task

We compared activity between localization and navigation phases of the MSN task to test whether sensorimotor and mnemonic regions are differentially engaged across task demands. We hypothesized that sensorimotor regions of interest (ROI), specifically the left primary motor cortex (M1), supplementary motor area (SMA), and right anterior cerebellum (CBL), would show increased activity during localization, where participants performed isometric exertions with their right hand. In contrast, we expected mnemonic regions, including the retrosplenial cortex (RSC), hippocampus (HC), and entorhinal cortex (ERC), to show greater activity during navigation, which required navigating the learned force-time space.

Sensorimotor regions showed stronger responses during localization than navigation (**Fig. 2E,F)**. Mean beta values for the navigation - localization contrast were negative in M1 (-1.26 ± 0.13), CBL (-0.45 ± 0.073), and SMA (-0.67 ± 0.10) and were all significantly less than zero (all p < 0.001 family-wise error (FEW) corrected over sensorimotor ROIs, **Fig. 2F**). Conversely, mnemonic regions were preferentially engaged during navigation (**Fig. 2G,H)**. Mean beta values for the navigation-localization contrast were positive in ERC (0.11 ± 0.060), HC (0.15 ± 0.050), and RSC (0.47 ± 0.10) and these activations were significantly greater than zero (all p < 0.05 FWE corrected over mnemonic ROIs, **Fig. 2H**). Together, these results demonstrate that the localization and navigation segments differentially engaged mnemonic and sensorimotor areas, enabling us to examine how each system represents task structure during the MSN task.

### The entorhinal cortex utilizes grid-like codes to represent sensorimotor information

We tested whether the ERC displays grid-like representations as participants navigated the force-time space using established fMRI methods^13–15^. Grid cells in the ERC spatially represent the environment through receptive fields arranged in a regular, hexagonal pattern, where, with respect to the grid orientation (ϕ), one can either traverse aligned (θa) or misaligned (θm) trajectories (**Fig. 3A**). Aligned trajectories will cross through more receptive fields and thus evoke greater grid cell activity than misaligned trajectories. Since nearby grid cells share similar orientation^18^, the ERC population activity is thought to display hexadirectional firing and can be modeled using a 6-fold cosine waveform: cos(6(*θ* - ϕ))^34^ (**Fig. 3B, C**).

**Figure 3:**
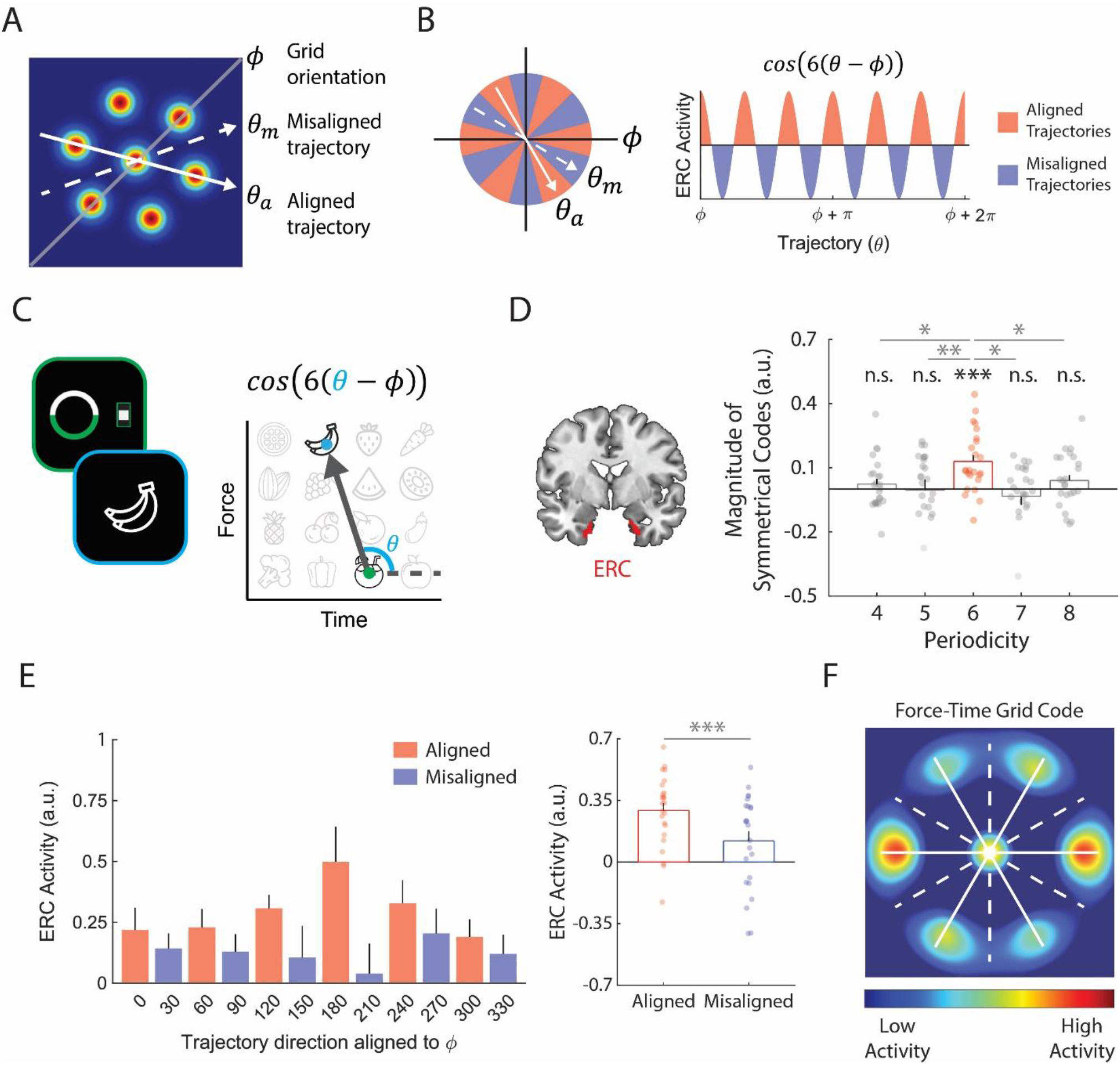
The entorhinal cortex utilizes grid-like codes to represent sensorimotor information. **(A)** Receptive fields of grid cells in the entorhinal cortex (ERC) are arranged hexagonally, resulting in different activity depending on the trajectory. Trajectories are categorized as aligned (θa) or misaligned (θm) with the grid orientation (ϕ). **(B)** ERC activity as a function of trajectory can be modeled as a 6-fold cosine waveform where aligned trajectories pass through more receptive fields and evoke greater activity than misaligned trajectories. **(C)** During the MSN task, the inferred trajectory, for each trial, was computed and adjusted by each participant’s estimated grid orientation as they traveled from the localized cue (green) to the target cue (blue) in the navigation segment. **(D)** ERC (anatomical mask in red) activity was significantly correlated with 6-fold modulation (*** p < 0.001; two-tailed one-sample Wilcoxon signed-rank test, corrected for number of periodicities (n = 5) using the Holm-Bonferroni method), but not with control periodicities (4,5,7,8 fold, all P > 0.05). Six-fold modulation signals in the ERC were also significantly greater than those of control periodicities (** p < 0.01, *p < 0.05; two-tailed paired Wilcoxon signed-rank test, corrected for number of comparisons (n = 4) using Holm-Bonferroni method). Error bars represent one sem. **(E)** Mean ERC activity was plotted separately for trajectories divided into 12 equal bins of 30° (Left) based on the direction of inferred trajectories (θ, C). The mean ERC activity for aligned trajectories was significantly greater than the mean ERC activity for misaligned trajectories (Right, *** p < 0.001, two - tailed paired t-test). Error bars represent one sem. **(F)** By interpolating ERC activity for aligned and misaligned trajectories in polar coordinates, one can conceptualize a force-time grid code where we observe a hexagonal firing modulation in ERC as participants navigated the force-time space.

We first estimated the putative grid orientation angle (ϕ) for each participant by decomposing the as regressors for cosine waveform into cos6 *θ* * cos6*ϕ* + sin6 *θ* * sin6*ϕ* and inputting cos6 *θ* and sin6 *θ* as regressors for predicting ERC activity using a general linear model (**GLM 1**, see Methods). We then utilized the beta values for cos6*θ* and sin6*θ* to estimate ϕ and tested for 6-fold periodicity signals in the ERC by inputting cos(6(*θ*-ϕ)) as a regressor in a second GLM (**GLM 2**, see Methods). We found significant non-uniform clustering of putative grid orientations (*ϕ*) for most participants (18/24) across voxels in the ERC (mean z **±** SE = 5.37 ± 0.80, **Supplementary Fig. 7**). To rule out selection bias, we implemented an unbiased cross-validation (CV) procedure where fMRI sessions were split into a ‘training’ and ‘testing’ datasets, estimates from the training dataset acquired in GLM 1 were used to test periodicity signals in the and *ϕ* testing dataset in GLM 2 (see **Methods**).

We found patterns of activity in the ERC to be significantly correlated with 6-fold periodicity (P < 0.001, FWE corrected) (**Fig. 3D**). Importantly, the ERC was not significantly explained by other control periodicities (n = 4,5,7,8, p > 0.05) and the 6-fold signal in the ERC was significantly greater than the control periodicities (all p < 0.01, FWE corrected) (**Figure 3D**). To further visualize this effect, we grouped mean ERC activity separately for aligned and misaligned trajectories (**Fig. 3E**, Left), for which we saw mean ERC activity for aligned trajectories to be significantly greater than misaligned trajectories (P < 0.001, **Fig. 3F**). Interpolating these results in polar coordinates revealed a hexagonal modulation pattern in ERC activity (**Fig. 3F**), resembling grid-like responses observed in the ERC of rodents during spatial navigation^7,16^. These results indicate that the grid-like properties of the ERC can extend to representing the sensorimotor domain.

We next examined whether hexadirectional signals aligned to the ERC’s grid orientation were present in other regions. No significant modulation was observed in sensorimotor regions (M1, CBL, SMA), or the hippocampus (all P > 0.05; **Supplementary Fig. 8A, B**). While we observed 6-fold modulation in the RSC, this effect did not survive correction for multiple comparisons across the number of tested ROIs (**Supplementary Fig. 8. A, B**). However, hexadirectional modulation strength was correlated between ERC and RSC across participants (Kendall’s τ = 0.36, p = 0.015, **Supplementary Fig. 9A**), suggesting potential functional coordination between these regions during the navigation of force-time space.

As further control, we independently estimated grid orientation for each region (excluding ERC) and assessed region-specific hexadirectional modulation (**Supplementary Fig. 8C**). We observed a significant 6-fold periodic signal in the SMA (**Supplementary Fig. 8D**). This response was selective to 6-fold modulation and not explained by control periodicities (all P > 0.05). However, hexadirectional activity in the SMA was not significantly correlated with ERC grid-like activity across participants (Kendall’s τ = 0.014, p = 0.94, **Supplementary Fig. 9B**), suggesting that this grid-like signal in the SMA may reflect an independent organization of sensorimotor representations^35^.

### Participants evaluate force and time differently when exerting and navigating sensorimotor space

During the MSN task, participants navigated a task-relevant force-time space in which force and time differences were instructed to be weighted equally. To test whether their behavior reflected such representation, we examined whether MSN accuracy was similarly influenced by spatial metrics derived separately from force and time dimensions (**Fig. 4A**) (**See Methods**). For each participant, MSN trial accuracy was modeled using logistic regression as a function of these spatial metrics, separately for force and time dimensions (**Fig. 4B**, bottom). Consistent with earlier results (**Fig. 2D**), all spatial metrics derived from force and time significantly predicted performance (all P < 0.05) and did not differ between dimensions (all P > 0.05), indicating participants were equally sensitive to force and time during navigation (**Fig. 4B**).

**Figure 4:**
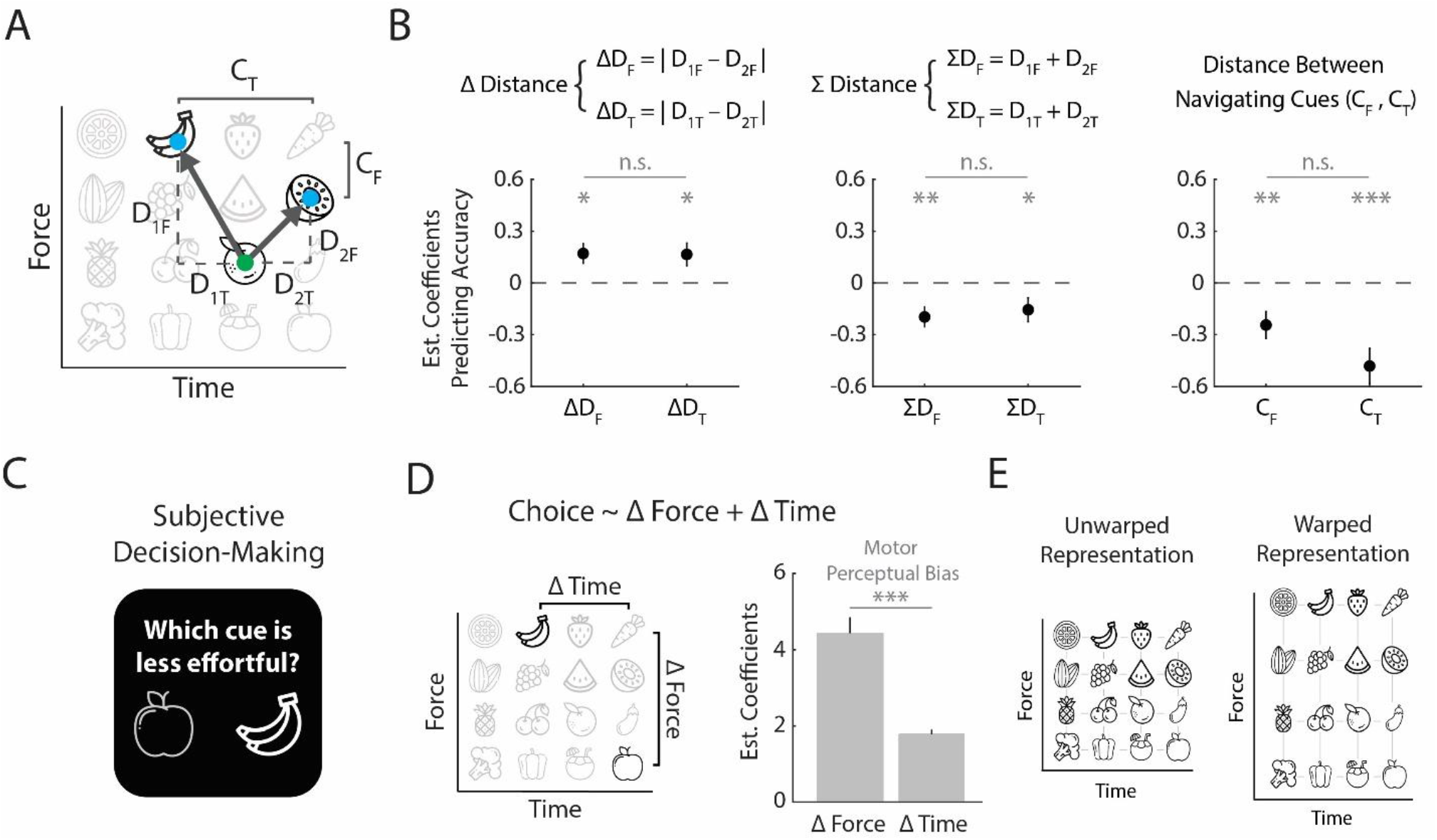
Participants differentially evaluate force and time when exerting and navigating sensorimotor space. **(A)** To investigate whether the spatially mediated metrics had different effects for force and time, Δ Distance and Σ Distance were computed separately for the force (ΔD_F_, ΣD_F_) and time (ΔD_T_, ΣD_T_) dimensions. Trajectory similarity was computed as the difference between the two cues in the navigation phase in the force (C_F_) and time (C_T_) levels. **(B)** A mixed effects generalized linear model was used to predict spatially mediated metrics separated in force and time dimensions for each participant. There was no significant difference between the spatially mediated metrics separated by force and time levels for Δ Distance, Σ Distance, and the distance between navigating cues (all p > 0.05, Wilcoxon paired signed-rank test). Error bars represent one sem. * p < 0.05, **p < 0.01, ***p < 0.001 **(C)** In the subjective decision-making task, participants were shown two cues and were instructed to select the cue that felt less effortful after considering differences in their force and time levels. **(D)** Participants’ responses in the subjective decision-making task were modeled as a linear combination of Δ Force and Δ Time, which were the differences in force and time levels in the force - time space (left). Participants weighted Δ Force significantly greater than Δ Time (***p < 0.001, Wilcoxon signed-rank test), which indicated a motor perceptual bias towards force levels. Error bars represent one sem. **(E)** Based on participants’ behavior in the MSN and subjective decision-making task, we hypothesized that while the MSN task required an unwarped, one-to-one representation of force-time space, a warped representation of force-time space in sensorimotor brain regions could reflect the motor-perceptual bias that individuals felt when performing the isometric exertions for different cues.

However, equivalent weighting during navigation did not imply equivalent subjective experience during the isometric exertions^30^. To examine how individuals subjectively perceived differences in force and time during exertions, participants completed a subjective decision -making task^30^ in which they selected the less effortful cue from a pair of cues (**Fig. 4C**). Using logistic regression, we modeled participants’ choices based on differences in force (Δ Force) and time (Δ Time) between the two cues (**Fig. 4D**). Across participants, Δ Force was perceived as significantly more effortful than Δ Time (paired t-test, p < 0.001) (**Fig. 4D**), revealing a robust motor perceptual bias towards force during exertion.

Together, these findings indicated that multiple force-time representations could be engaged during the MSN task. While performing the MSN task required a force-time space with equivalent representations of the force and time dimensions, their motor perceptual bias during exertions indicated potential existence of a warped force-time space with greater bias for force. This dissociation led us to hypothesize that different brain regions could encode distinct force-time spaces: sensorimotor regions responsible for generating motor commands may represent a warped force-time space (**Fig. 4E**; right), causing participants to perceive force differences as more effortful, whereas mnemonic regions could encode an unwarped force-time space (**Fig. 4E**; left), reflecting the learned associations participants made between cues and exertions.

### Sensorimotor and mnemonic brain regions differentially represent the force and time of exertion

To test whether sensorimotor and mnemonic regions encode distinct force-time representations, we utilized pattern component modeling (PCM)^36^ to assess the representational structure within each region. PCM utilizes covariance (second-moment) matrices as a central statistical quantity to describe the representational content of brain regions during different conditions^36^.

We focused our PCM analysis on the localization segment, during which participants actively performed isometric exertions and recalled the matching cue, which would engage both sensorimotor and mnemonic processes. We leveraged the eight most frequently sampled cues (**Fig. 5A**, yellow) to build force (f) and time (t) feature sets that define the sampled cues based on their respective force and time levels. These feature sets were used to compute predicted covariance matrices that describe the representational structure of each cue by differences in the force (**Fig. 5A**, left) and time (**Fig. 5A**, right) dimensions. We then evaluated how well the force and time covariance matrices explained differences in activity patterns within each ROI by comparing the cross-validated log-likelihood values of three different component models: A force only model (F), which modeled the neural covariance structure based on the force component, a time only model (T), which modeled the structure based on the time component, and a force + time model (F+T), which modeled the structure using a weighted linear combination of both force and time component matrices. To test whether sensorimotor and mnemonic brain regions differentially represent force and time, we computed the log Bayes factor (BF_F vs. T_), which is the difference in the log model evidence for the force and time only models (see **Methods**). In addition, we compared the force and time component weights from the F + T model to test for significant differences in weighting the two dimensions for each ROI.

**Figure 5:**
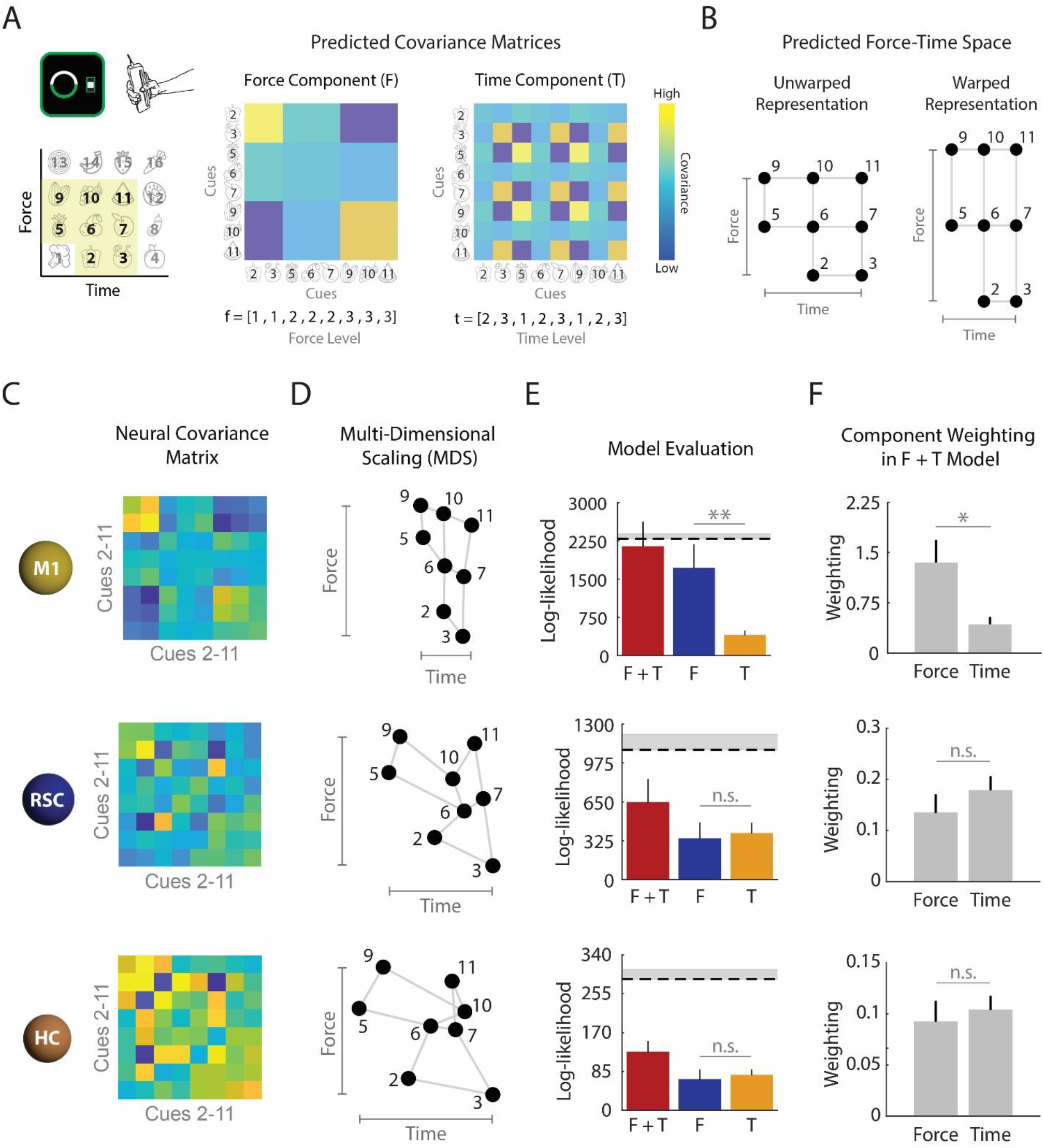
Sensorimotor and mnemonic regions differentially represent the force and time of exertions. **(A)** Pattern component modeling (PCM) analysis was performed to understand how different brain regions represent the force and time of exertion during the localization condition. The PCM analysis included exertion data from 8 cues (yellow box, numbered cues 2-11) as those cues were sampled most frequently in the MSN task. Force (f) and time (t) feature sets were computed that defined the cues based on their respective force and time levels. These feature sets were used to compute predicted covariance matrices that described the covariance of a pair of cues due to either differences in force or time levels. **(B)** Predicted force-time space from the sampled cues 2-11 that depicts an unwarped representation (left), where force and time features are equally represented, and a warped representation (right), where the cues are represented more saliently in force compared to time features. **(C)** PCM results for M1, RSC, and HC. The cross-validated estimates of the neural second moment matrix describe the activity profile distribution for each brain region during the localization segment where participants performed exertions corresponding to cues 2-11. **(D)** Multi-Dimensional Scaling was performed (middle column), where cues 2-11 were projected on the eigenvectors of the second moment matrix for each representative brain region. **(E)** The performance of each model (right column) for predicting the neural covariance matrix. The force (F) and time (T) models predicted the neural second moment matrices using both F and T models. The force and time (F + T) model utilized both components. The dotted line represents the lower bound noise ceiling. Error bars represent one sem. ** p < 0.01; two-tailed paired Wilcoxon signed-rank test, corrected for number of sensorimotor brain regions (n = 3) using Holm-Bonferroni method. **(F)** The weights associated with the force and time components in the F + T model. The force component was weighted significantly greater than the time component for M1. Error bars represent one sem. * p < 0.05; two-tailed paired Wilcoxon signed-rank test, corrected for number of sensorimotor brain regions (n = 3) using Holm-Bonferroni method.

We hypothesized that sensorimotor regions would exhibit a warped force-time space, where force features are more saliently represented than that of time (**Fig. 5B**, right). In line with our hypothesis, we found that sensorimotor regions such as the M1 and CBL encoded the force dimension more strongly than the time dimension. The covariance matrix for M1 and CBL activity resembled the force component more than the time component (**Fig. 5C**, **Supplementary Fig. 10A**), and multi-dimensional scaling (MDS) revealed a stretched representational space along the force axis, consistent with a warped force - time representation (**Fig. 5D**, **Supplementary Fig. 10B**). While the F + T model performed the best in both M1 and CBL, the force-only model performed similarly and outperformed the time-only model for both the M1 (mean BF_F vs. T_ ± SE: 1312 ± 476, p < 0.01, corrected for sensorimotor ROIs) (**Fig. 5E**) and the CBL (mean BF_F vs. T_ ± SE: 659 ± 304, p < 0.05, corrected for sensorimotor ROIs) (**Supplementary Fig. 10D**).

Furthermore, within the F + T model, the force component weight was significantly greater than the time component for both the M1 (p < 0.05, corrected for sensorimotor ROIs, **Fig. 5F**) and CBL (p < 0.05, corrected for sensorimotor ROIs, **Supplementary Fig. 10D**). These results confirmed a warped force - time space in M1 and CBL, with both regions showing a stronger representation of force than time.

In contrast, the SMA showed a more unwarped representation of force and time dimensions. The neural covariance matrix and MDS from activity patterns within the SMA revealed no clear dominance of either feature (**Supplementary Fig. 10A**, **B**). When comparing each model’s performance, the F + T model performed the best, with the F and T models performing comparably (mean BF_F vs. T_ ± SE: 34 ± 161, p > 0.05) (**Supplementary Fig. 10C**). There was no significant difference in the weighting assigned to the force and time components in the F + T model (**Supplementary Fig. 10D**). This pattern may reflect the role of SMA in higher-order movement aspects such as timing^18^ and movement inhibition^19– 21^, which were key components of the exertion-release behavioral structure in the localization segment of the MSN task.

Mnemonic regions exhibited similarly unwarped representations. The neural covariance matrix for RSC, HC and ERC showed comparable sensitivity to force and time (**Fig. 5C–D; Supplementary Fig. 10**). Additionally, model comparisons indicated similar performance of the F and T models for the RSC (mean BF_F vs. T_ ± SE:-43 ± 123, p > 0.05), HC (mean BF_F vs. T_ ± SE:-9.6 ± 25, p > 0.05), and ERC (mean BF_F vs. T_ ± SE: - 0.3 ± 4, p > 0.05) and component weights in the F + T model did not differ between force and time (all p > 0.05). (**Fig. 5E, F**, **Supplementary Fig. 10C, D**). Together, these results indicate that while sensorimotor regions encode a force-biased, warped force-time space, mnemonic regions preserve an unwarped, task-relevant representation of force and time during the MSN task.

### Force and time information is flexibly transformed between sensorimotor and mnemonic regions

The PCM results suggested that force-time information may be transformed as it flows from sensorimotor regions, which encode a force-biased space, to mnemonic regions, which preserve an unwarped, task-relevant representation. We therefore tested whether the functional coupling between these systems supports such transformation. Given its anatomical and functional connectivity with both regions, we initially proposed the RSC to act as a key intermediary in facilitating this transformation^20– 22,37^.

We first used a psychophysiological interaction (PPI) analysis with the RSC as the seed, analyzing the interaction between RSC activity and regressors from the localization–navigation contrast (**Fig. 2E-H**). Consistent with our initial hypothesis, we found that the RSC had significantly greater coupling with M1, SMA, and HC during the localization phase (**Supplementary Fig. 11**, p < 0.05, corrected for the number of ROIs (n = 5)). These findings provide initial evidence that the RSC can dynamically coordinate with both sensorimotor and mnemonic regions as participants associate isometric exertions with their corresponding contextual cues.

Building on the observed functional coupling, we applied dynamic causal modeling (DCM) to examine directed causal interactions between the sensorimotor and mnemonic brain regions. We specified a deterministic, bilinear DCM to test directed influences between the RSC, M1, SMA, and HC (i.e., effective connectivity), which we chose as seed regions based on our PPI findings. We employed the Parametric Empirical Bayes (PEB) framework^38,39,^ estimating a single ‘full’ model that included all relevant connections for each participant (1^st^ level), then examining commonalities and differences in connection parameters across participants (2^nd^ level) (SI – Dynamic Causal Modeling).

Given the large number of connection parameters, we interpreted connections with posterior probability greater than 0.99, which indicated very strong evidence (**Supplementary Fig. 12B, C**). During localization, we identified extensive modulatory connections between mnemonic and sensorimotor regions (**Supplementary Fig. 12C; Supplementary Table 2**). Consistent with our PPI findings, the RSC inputs from M1, SMA, and HC. Notably, the nature of these connections varied by region type: connections originating from sensorimotor regions were consistently inhibitory, while those from mnemonic regions were excitatory (**Supplementary Fig. 12C**). These findings potentially indicated directionally specific interactions between sensorimotor and mnemonic regions as participants associated exertions with learned cues.

### Inhibitory connectivity from M1 determines the degree of force-warping in mnemonic force-time space

Among the modulatory connections, the inhibitory influence from M1 to RSC was particularly relevant for potentially transforming force-time representations (**Supplementary Fig. 12C**, **Fig. 6A**). Because M1 encoded a force-biased space, we tested whether this inhibitory input reduced force weighting in RSC, resulting in a less warped force-time space (**Fig. 6A**). For each participant, we quantified the weighting of force (ω_F_) and time (ω_T_) components in the F + T model from the PCM analysis (**Fig. 5F**, **6A**). We defined the downregulation of force in RSC as the normalized difference between ω_F_ and ω_T_ ((ω_T_ - ω_F_) /(ω_T_ + ω_F_)), where values closer to 1 indicated reduced force weighting whereas values closer to - 1 indicated increased force weighting relative to time. This index was then used as a second-level covariate in our PEB analysis to model individual differences in the M1-to-RSC connection parameter.

**Figure 6:**
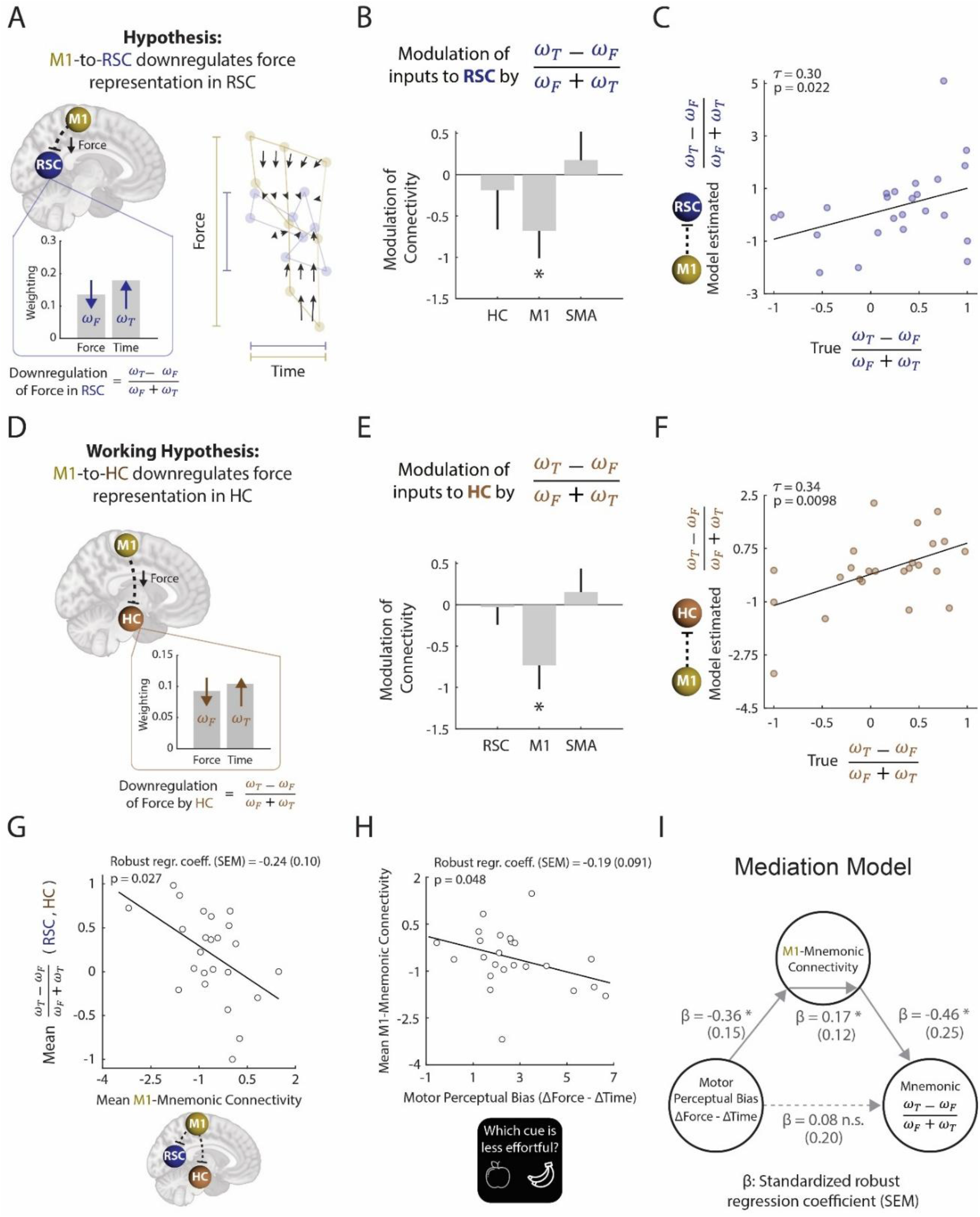
Inhibitory connectivity from M1 determines the degree of force-warping in mnemonic force-time space. **(A)** Using dynamic causal modeling (DCM), we identified very strong evidence (posterior probability > 0.99) of inhibitory modulation from M1 to RSC at the time of localization. We hypothesized that this inhibitory modulation from M1 to RSC could downregulate force representation in the RSC, which would reflect the transformation from a warped to a balanced force-time space. To test this hypothesis, we asked whether individuals with stronger inhibitory modulation from M1 to RSC showed a greater downregulation of force weighting: (ω_T_ – ω_F_)/ (ω_F_ + ω_T_), where ω_F_ and ω_T_ were the force and time weighting metrics in RSC computed from the F + T component model in the PCM analysis. **(B)** We assigned the downregulation of force ((ω_T_ – ω_F_)/ (ω_F_ + ω_T_)) in RSC as a second-level covariate in the original DCM analysis using the parametric empirical bayes (PEB) framework. We found that there was very strong evidence (* posterior probability > 0.99) of force downregulation in RSC modulating M1-to-RSC connectivity. Specifically, those who exhibited stronger downregulation of force in RSC showed greater negative modulation from M1 to RSC. Error bars represent 90% credible interval. **(C)** Using leave-one-out cross-validation, we computed out-of-sample estimation of participants’ downregulation of force in RSC using the degree of inhibition of the RSC by M1 (y-axis) and compared it to the true downregulation of force for that participant’s RSC. **(D)** DCM additionally revealed very strong evidence of inhibitory modulation (posterior probability > 0.99) from M1 to HC at time of localization. Similar to RSC, we hypothesized that inhibitory modulation from M1 to HC could downregulate force representation in the HC. **(E)** There was very strong evidence (* posterior probability > 0.99) of downregulation of force in HC modulating M1-to-HC connectivity. Error bars represent 90% credible interval. **(F)** Using leave-one-out cross-validation, we computed out-of-sample estimation of participants’ downregulation of force in HC using the degree of inhibition of the HC by M1 (y-axis) and compared it to the true downregulation of force for that participant’s HC. **(G)** Participants with greater negative mean connectivity (stronger inhibition) from M1 to mnemonic areas (RSC, HC) had increased mean downregulation of force in mnemonic areas (RSC, HC). **(H)** Participants that showed greater motor perceptual bias in the subjective decision-making task showed more negative mean connectivity (stronger inhibition) from M1 to mnemonic areas (RSC, HC). **(I)** A mediation analysis revealed that M1-to-Mnemonic connectivity fully mediated the relationship between motor perceptual bias and downregulation of force in mnemonic regions. * p < 0.05.

We found very strong evidence (posterior probability > 0.99) that the strength of the inhibitory M1 - to-RSC connection was associated with increased downregulation of force in RSC. Thus, participants with reduced force representation in RSC received stronger inhibition from M1 (**Fig. 6B**). To validate this relationship, we performed a leave-one-out (LOO) cross-validation analysis, where the PEB model was refitted excluding one participant and the M1-to-RSC connection parameters were used to predict the left-out participants’ force downregulation in RSC. The predicted and actual estimates were significantly correlated across participants (Kendall’s τ = 0.30, p = 0.022, **Fig. 6C**), providing confirmation that inhibitory M1-to-RSC connection could modulate the transformation of force-time representations.

Since M1 also showed inhibitory projections to the HC during localization (**Supplementary Fig. 12C**), we repeated the analysis and the same pattern was observed where stronger M1-to-HC inhibition predicted greater force downregulation in HC (**Fig. 6D-F**). When averaging across both RSC and HC, the mean M1-Mnemonic connectivity significantly predicted mean force downregulation (β(SE) = - 0.24(0.10), P = 0.027) (**Fig. 6G**) in RSC and HC, indicating that M1 broadly influences the transformation of force-time space across mnemonic areas.

Next, we asked whether individual differences in behavior shaped this transformation. Participants’ motor perceptual bias towards force during the subjective decision-making task (Δ Force - Δ Time, **Fig. 4C, D**) significantly predicted mean M1-Mnemonic inhibitory connectivity (β(SE) = -0.19(0.091), p = 0.048) (**Fig. 6H**), indicating that individuals with greater motor perceptual bias exhibited greater suppression from M1 to mnemonic regions. We then performed a mediation analysis to test whether participants’ motor perceptual bias (independent variable) affected the force downregulation in mnemonic regions (dependent variable) through the M1-Mnemonic connectivity (mediator). We found a significant indirect effect (ab (SE) = 0.17 (0.12), p = 0.047) (**Fig. 6I**), which indicated that greater motor perceptual bias towards force during exertions led to greater inhibition from M1 to mnemonic regions (a (SE) = -0.36 (0.15), p = 0.018), which in turn predicted greater downregulation of force in the mnemonic regions (b (SE) = -0.46 (0.25), p = 0.023). Importantly, the direct effect was not significant (c’ (SE) = 0.08, p = 0.51), which indicated that M1-mnemonic connectivity fully mediated how motor perceptual bias influenced the force-time representation in mnemonic brain regions. Together, these findings demonstrate that individual differences in motor perceptual bias shapes how force-time information is causally transformed between sensorimotor and mnemonic regions, reflecting a transition from an effort-based space to a task-relevant force-time space.

### The warping of force-time space in mnemonic regions is influenced by learning rate and motor perceptual bias

As cognitive maps can be shaped by experiences^40–43^, we next investigated whether individual differences in learning the cue-exertion pairs or subjective effort influenced how the force-time space was represented in mnemonic regions. We modeled individual participants’ learning curves across sessions using an exponential function: 1 – 0.5e^-bs^, where b represents the learning rate and s is the session number (**Supplementary Fig. 13**). Fast learners (blue) showed higher b values (e.g., b = 1.1) (**Fig. 7A**) and improved quickly while maintaining high performance in later sessions. In contrast, slow learners (red) had lower b values (e.g., b = 0.18) (**Fig. 7A**) and gradually improved in the MSN task, reaching high accuracy in later sessions.

**Figure 7:**
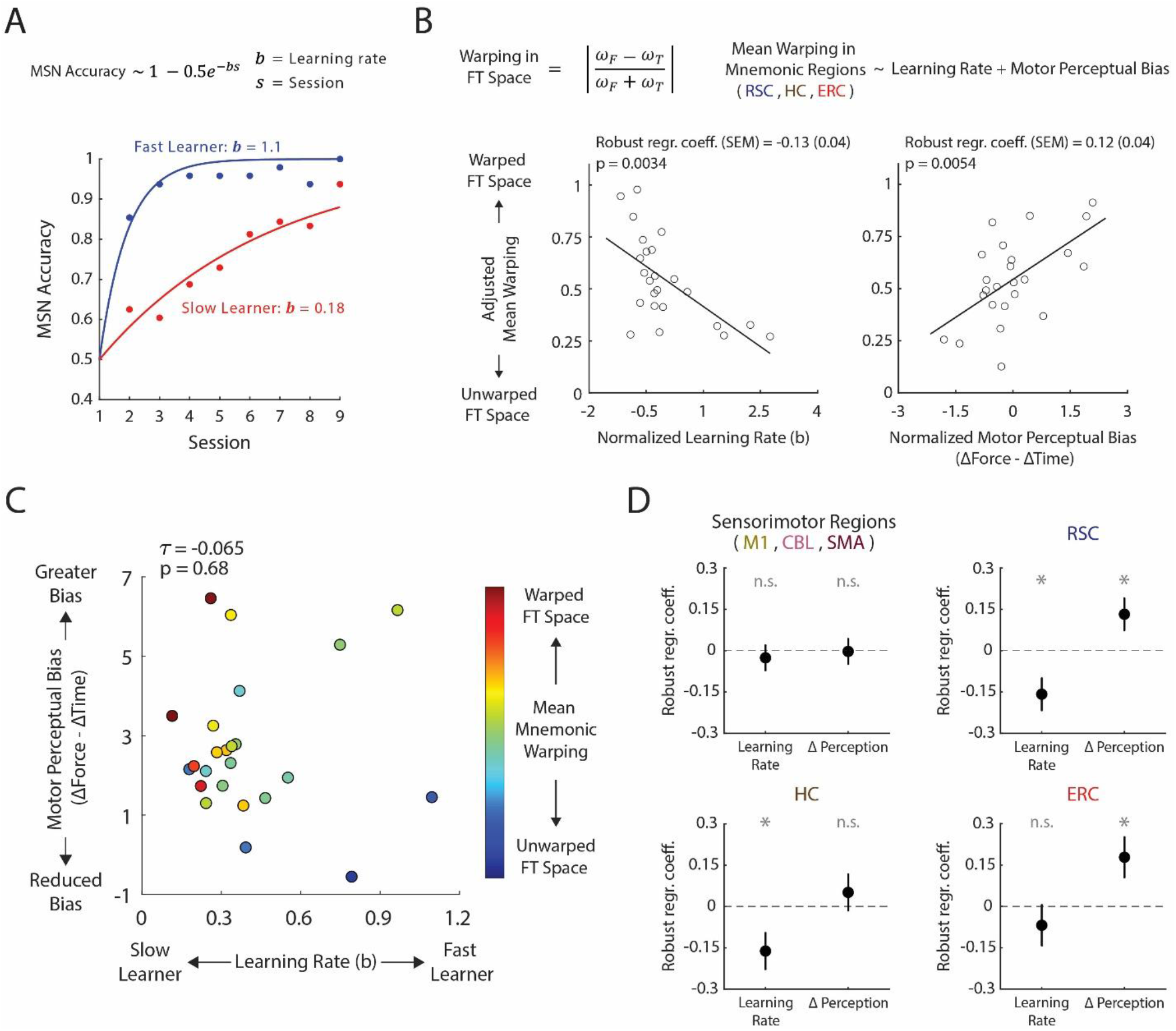
The warping of force-time space in mnemonic regions is influenced by learning and motor perceptual bias. **(A)** To investigate whether learning rate influenced individuals’ representation of force-time space in the RSC, we first captured the learning rate of individuals in the MSN task using an exponential function, where fast learners (representative participant in green, higher b) stabilized at a high performance level and slow learners (representative participant in red, lower b) gradually improved their performance. **(B)** We defined a warping metric of force-time space in mnemonic regions by computing the absolute value of (ω_F_ - ω_T_)/(ω_T_ + ω_F_) for each participant. Values closer to 1 indicated a warped force-time space, while values closer to 0 indicated a more balanced force-time space. We found that individuals with faster learning rates exhibited a more balanced force-time space in the RSC, which indicated a more stable representation of force and time. Robust linear regression was used to model mean warping in all mnemonic regions (RSC, HC, ERC) as a function of learning rate and motor perceptual bias. Both learning rate and motor perceptual bias were significant predictors of warping in mnemonic regions. **(C)** Visualization of participants based on their learning rate and motor perceptual bias, with color indicating the mean warping in mnemonic brain regions. Faster learners exhibited a less warped, more unbiased force-time space (blue) whereas individuals with a greater perceptual bias exhibited a more warped force-time space (red). **(D)** Linear regressions were used to separately model mean warping in sensorimotor regions (top left), RSC (top right), HC (bottom left), and ERC (bottom right) as a function of learning rate and motor perceptual bias. * p < 0.05

We hypothesized that fast learners would exhibit a less warped force-time space in mnemonic regions, having had more time to consolidate the task-relevant space. In addition, based on our connectivity results (**Fig. 6G-I**), we hypothesized that individuals with stronger motor perceptual bias toward force would show greater warping of force-time space in mnemonic regions. To test these hypotheses, we computed a warping metric (**Fig. 7B**), which unlike the previously defined downregulation of force (**Fig. 6A**), involved taking the absolute normalized difference between ω_F_ and ω_T_ (|(ω_T_ - ω_F_) /(ω_T_ + ω_F_)|), where values closer to 0 indicated an unwarped representation of force and time, and values closer to 1 indicating warping of force-time space.

A robust linear regression was used to model mean warping in all three mnemonic regions (RSC, HC, ERC) as a function of learning rate (b, **Fig. 7A**) and motor perceptual bias (Δ Force - Δ Time, **Fig. 4C, D**). Consistent with our hypotheses, individuals with higher learning rates exhibited significantly less warped force-time representations in mnemonic regions (β(SE) = -0.13(0.04), p = 0.0034) (**Fig. 7B**, **Supplementary Fig. 14A**), suggesting that earlier acquisition of the contextual force-time space led to less warped representation of force and time in mnemonic regions. In contrast, motor perceptual bias was a significant positive predictor of warping in mnemonic regions (β(SE) = 0.12(0.04), p = 0.0054) (**Fig. 7B**, **Supplementary Fig. 14A**), confirming that participants who perceive force as more effortful exhibited a more warped force-time space in mnemonic regions. Importantly, learning rate and motor perceptual bias were not significantly correlated (Kendall’s τ = -0.065, p = 0.68, **Fig. 7C**), which suggested that these behavioral factors independently influenced the warping of the force-time space in mnemonic regions. Mean warping in sensorimotor brain regions was not significantly predicted by either learning rate (β(SE) = -0.03(0.05),p = 0.58) or motor perceptual bias (β(SE) = 0.00 (0.05), p = 0.95), indicating that these behavioral factors selectively influenced representations in mnemonic but not sensorimotor regions (**Fig. 7D**, **Supplementary Fig. 14B**).

Within mnemonic regions, we observed different patterns of sensitivity to the behavioral metrics (**Fig. 7D**). Warping in RSC was significantly associated with both learning rate (β(SE) = -0.16(0.06), p = 0.01) and motor perceptual bias (β(SE) = 0.13(0.06), p = 0.04), consistent with its proposed role in integrating mnemonic and sensorimotor processes^20–22,37^ (**Supplementary Fig. 14C**). In the HC, warping was significantly associated with learning rate (β(SE) = -0.16(0.07), p = 0.02) but not motor perceptual bias (β(SE) = 0.05(0.07), p = 0.45), suggesting its primary role in consolidating the learned, unwarped force-time structure^41,44–46^ (**Supplementary Fig. 14D**). In contrast, warping in the ERC was significantly associated with motor perceptual bias (β(SE) = 0.18(0.07), p = 0.03), but not learning rate (β(SE) = - 0.07(0.07),p = 0.37), consistent with prior work showing that distortions in perceptual or environmental inputs can alter the spatial coding properties of the ERC^42,47^ (**Supplementary Fig. 14E**).

### The HC and RSC reflect distinct computational processes when navigating sensorimotor space

Given that the representational structure of force-time space in the HC and RSC was shaped by learning rate, we next asked whether these mnemonic regions supported distinct computational processes during the MSN task. We hypothesized that the HC, given its established role in path integration^48–50^, would reflect individual differences in sensitivity to spatial metrics during mental traversal of the force-time space (**Fig. 8A**). We computed the bias for each spatial metric (**Fig. 4B**), defined as the difference between parameters derived from the force and time dimensions (e.g., Δ Distance Bias = ΔD_T_ - ΔD_F_). We then examined whether these bias metrics were associated with ω_F_ and ω_T_ in mnemonic regions. Notably, Σ Distance Bias was significantly correlated with the downregulation of force in the HC (Kendall’s τ = 0.42, p = 0.042, FWE corrected over number of correlations (n = 9)) (**Supplementary Fig. 15A**). To test whether this effect was specific to the HC, we conducted a robust linear regression where Σ Distance Bias was modeled as a function of force downregulation in RSC, HC and the ERC. Consistent with our hypothesis, only force downregulation in the HC significantly predicted Σ Distance Bias (β(SE) = 0.30(0.11), p = 0.01) (**Fig. 8B**, **Supplementary Fig. 15B**), indicating that individuals who showed reduced force representation relative to time in HC were also more sensitive to errors that increased with greater traversal distance in the force relative to time dimension (**Fig. 8C**).

**Figure 8:**
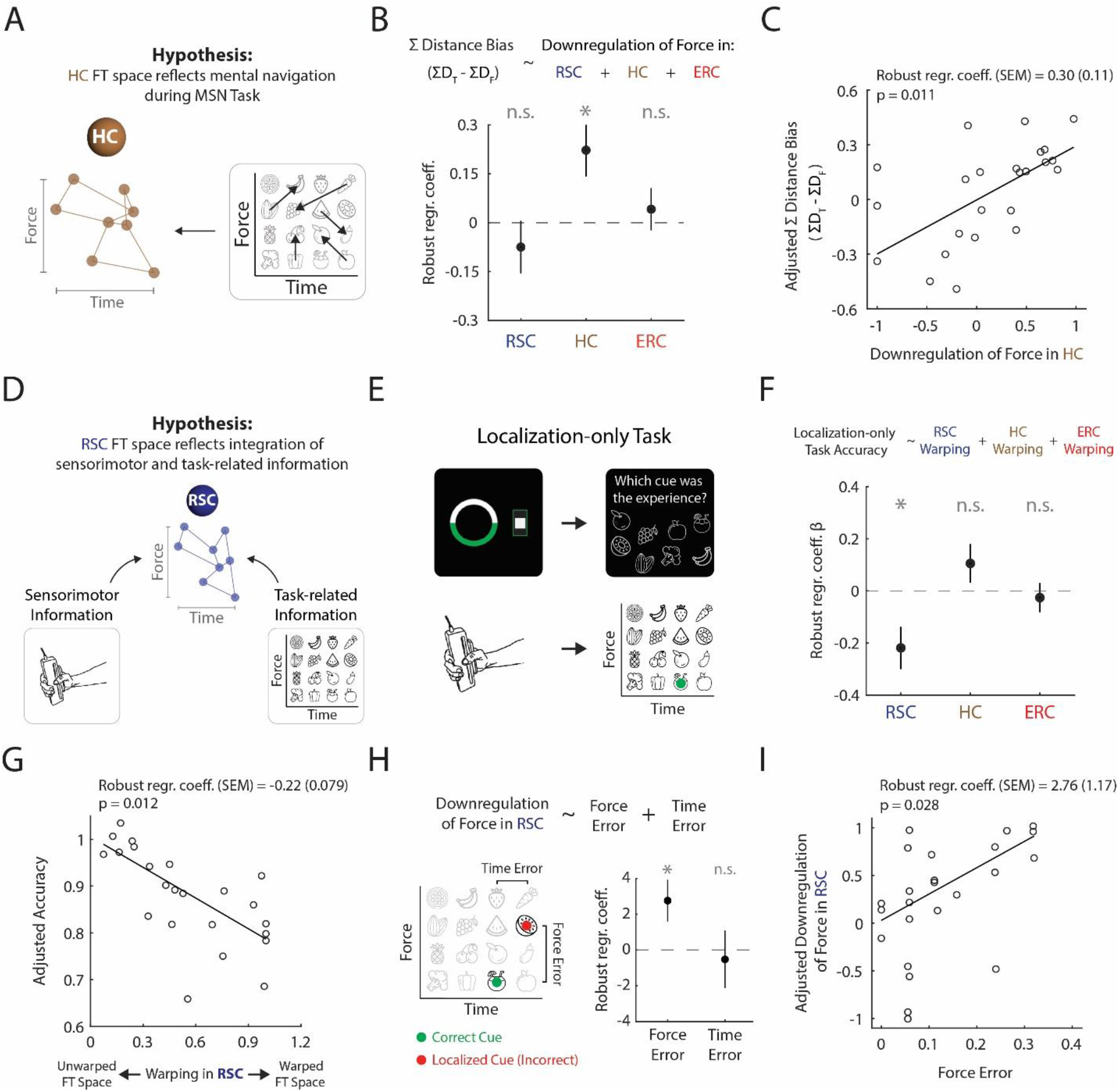
The HC and RSC reflect distinct computational processes when navigating sensorimotor space. **(A)** Given the role of HC in path integration during spatial navigation, we hypothesized that interindividual differences in the force-time space in HC should reflect the sensitivity of participants to spatially mediated metrics during the MSN task. **(B)** Robust linear regression was used to model Σ Distance Bias, quantified as the difference in sensitivity to the total distance traveled in the time and force dimensions (ΣD_T_ – ΣD_F_) using the downregulation of force in RSC, HC and ERC. The downregulation of force in HC significantly predicted Σ Distance Bias. Error bars represent one sem. * p < 0.05 **(C)** Visualizing the effect in (B), participants that exhibited greater downregulation of force in HC were more prone to errors in the MSN task with increase in distance in the force dimension (more positive ΣD_T_ – ΣD_F_). **(D)** Given the role of RSC as a hub that is interconnected with sensorimotor and mnemonic processes, we hypothesized that the force-time space in RSC should reflect participants’ ability to accurately associate exertions with its matching cue. **(E)** To assess participants’ ability to accurately associate exertions with their matching cue, participants performed a localization-only task at the end of Day 3 where they performed an exertion identical to the localization segment in the MSN task. They were then presented with a screen showing a correct cue that matched the exertion profile in force and time level as well as seven other distractor cues. **(F)** Participants’ performance in the localization-only task was modeled as a function of warping metrics in the RSC, HC and ERC. Warping in RSC was a significant predictor of participants’ accuracy in the localization-only task. Error bars represent one sem. * p < 0.05 **(G)** Visualizing the effect in (F), participants that exhibited a more warped force-time space in the RSC showed reduced accuracy in the localization-only task. **(H)** The downregulation of force in RSC was modeled as a function of force and time error in the localization-only task. Force and time errors were defined as the average distance between the correct and localized cue in force and time dimension respectively across all trials. Force error was a significant predictor of force downregulation in the RSC. Error bars represent one sem. * p < 0.05 **(I)** Visualizing the effect in (H), participants that made more errors in the force dimension exhibited greater downregulation of force in the RSC.

We next tested whether the RSC, given its functional and anatomical coupling with both sensorimotor and mnemonic systems^20–22,37^, integrated both sensorimotor and task-related information (**Fig. 8D**). We first computed each participant’s performance in the localization-only task (**Fig. 8E**, **Supplementary Fig. 5**) as their accuracy was indicative of how well participants integrated sensorimotor feedback with the learned cues in the force-time space. We then modeled participants’ performance in the localization-only task as a function of the warping metric (**Fig. 7B**) in each mnemonic region. Warping in the RSC was a significant negative predictor of participants’ accuracy in the localization-only task (β(SE) = -0.22(0.08), p = 0.01) (**Fig. 8F**, **Supplementary Fig. 15C**), indicating that greater distortion of force-time space in the RSC was associated with increased localization errors (**Fig. 8G**).

Lastly, we asked whether the errors in the localization-only task were specifically related to how force and time were represented in the RSC. We modeled force downregulation in the RSC as a function of errors made along the force and time dimensions during the localization-only task, where errors were quantified as the distance between the correct and chosen cue in each dimension (**Fig. 8H**). Force error was a significant positive predictor of force downregulation in the RSC (β(SE) = 2.76(1.17),p = 0.03) (**Fig. 8H**, **Supplementary Fig. 15D**), indicating that participants who made more errors in the force dimension exhibited reduced representation of force relative to time in the RSC (**Fig. 8I**). All together, these results indicate that the HC and RSC support distinct but complementary computational roles when representing sensorimotor space. While the HC reflects the navigation of force-time space, the RSC integrates sensorimotor feedback to support accurate localization in the learned force-time space.

## Discussion

Cognition and motor control are tightly interconnected^11,12,37^, yet the neural mechanisms by which the brain links movements with specific exogenous cues or task demands remains poorly understood. Cognitive map theory proposes that the brain represents relevant structured information to guide adaptive behavior^14,15,34,51,52^, but how such maps extend to the organization of endogenous sensorimotor memories has remained unclear. Our findings demonstrate that sensorimotor and mnemonic regions form distinct sensorimotor representations that dynamically communicate to encode learned associations between endogenous movement features and exogenous cues. Our findings integrate principles from motor control and cognitive mapping, providing a mechanistic account of how cognitive maps can be flexibly transformed to facilitate sensorimotor coordination during contextual control.

We provide direct evidence that ERC grid-like codes represent a cognitive map linking endogenous sensorimotor features with exogenous task-relevant visual cues. ERC activity exhibited hexadirectional coding for inferred trajectories in force-time space. Our finding aligns with prior studies showing how grid-like encoding in ERC extends beyond physical navigation to represent abstract spaces^26,29^, and expands this framework to endogenous abstract spaces including sensorimotor domains. Our results complement a recent finding reporting grid-like representations of action plans in the ERC^53^, which required participants to imagine different motor actions. In contrast, in our MSN task, participants first performed isometric exertions and then incorporated the sensorimotor feedback to navigate the learned force-time space. Thus, ERC grid codes can organize sensorimotor dimensions derived from both planned and experienced actions, supporting a cognitive map that links movement execution and endogenous sensorimotor features with exogenous task-relevant cues.

Our findings reveal grid-like encodings in other cortical regions, providing further insight into a broader network that could support cognitive maps for sensorimotor control. We observed evidence of grid-like responses in the RSC that were aligned to ERC’s grid orientation. The RSC is thought to play a key role in spatial navigation and mnemonic representation, storing and retrieving relevant spatial and contextual representations^20,54^. Prior studies have demonstrated grid-like RSC activity in humans undergoing navigation tasks during fMRI^26,29^ and direct electrophysiological recordings^55^. In addition, spatially periodic RSC signals have also been shown in rodents^23^. Interestingly, we found that participants who exhibited stronger ERC grid-like encoding also showed more robust grid-like signals in the RSC, supporting the idea of synchronized grid-like modulation across distributed brain regions^26,29,34,53,56^. While we also detected grid-like responses in the SMA, the signals were not aligned to the ERC’s grid orientation, and the grid-like strength was not correlated with that of the ERC, suggesting that the SMA may host an independent grid-like code. Given SMA’s established role in higher-order motor processes such as action planning^8^ and mental simulation^53^, both of which were crucial for solving the MSN paradigm, our results suggest that the SMA may utilize spatial mapping to organize sensorimotor information. Indeed, a recent study found grid-like firing in the rodent somatosensory cortex^35^. Together, these results suggest that grid-like representations may extend beyond mnemonic areas to include somatosensory regions, supporting the spatial organization of diverse forms of task - relevant information.

Our findings offer further insight into how endogenous movement information can be differentially represented across sensorimotor brain regions. We observe that the M1 and CBL show a significant bias toward the force over time of motor execution, resulting in a warped force-time space. The warping of force-time space mirrors participants’ robust motor perceptual bias, in which changes in force were perceived as more effortful than changes in time levels. Our findings suggest that M1 and CBL may encode an endogenous effort-based force-time space, reflecting the greater, neuromuscular demands associated with increasing force levels compared to prolonging exertion duration. Consistent with this interpretation, prior work has shown strong relationships between BOLD activity in M1 and CBL and electromyographic (EMG) signals during isometric exertions^57^. Additionally, repetitive transcranial magnetic stimulation (rTMS) applied to M1 can alter the subjective sense of effort^6^. In contrast to M1 and CBL, the SMA displayed a more balanced representation of force and time. Given SMA’s well-established role in higher-order movement processes such as planning^58^, timing^59^, and inhibition^9,60^, this region may prioritize temporal aspects of exertions. Together, these results highlight how distinct sensorimotor regions emphasize different dimensions of the force-time space, reflecting their specialized roles in generating, timing, and evaluation of endogenous effortful actions.

We found that mnemonic regions encoded an unwarped force–time space that reflected the task - relevant associations participants formed between visual cues and their sensorimotor features. During the MSN task, participants learned to weigh differences in force and time levels equally, and this unwarped mapping was preserved in mnemonic brain regions. This finding aligns with prior work showing that mnemonic regions represent relational structures across diverse exogenous domains, including value^61,62^, concepts^29,63^, and social hierarchies^26^, and extends this framework to the organization of sensorimotor features. Importantly, whereas M1 and the CBL encoded an effort-weighted force–time space driven by motor perceptual bias, the RSC, ERC and HC maintained a task-relevant force–time space in which force and time were weighted equally, consistent with the task-specific rules participants needed to follow in the MSN task. Our results suggest that mnemonic regions can store structured representations of visual cues according to their exogenous task-relevant associations^64^, even when those representations differ from the endogenous perceptual biases experienced during movement execution.

Our connectivity analyses provide mechanistic insight into how endogenous sensorimotor signals reflecting an individual’s internal state can be transformed into an exogenous task-relevant cognitive map. We show that inhibitory modulations from M1 to the RSC and HC mediate the degree to which force representations are downregulated in these regions. Participants with stronger inhibitory coupling from M1 exhibited a greater reduction of force weighting in mnemonic regions, transforming an effort-based force-time space to an unwarped task-relevant force-time space. Importantly, mediation analysis further revealed that individual differences in motor perceptual bias shaped mnemonic representations indirectly through M1-mnemonic coupling. These findings extend a growing body of work showing that the geometry of task-relevant representations can flexibly change in prefrontal and visual networks^27,65^. Here, we provide novel evidence that similar transformations in representational geometry also occur across networks: first, by demonstrating that representational space is reshaped as information flows from sensorimotor to mnemonic regions, and second, by showing that functional coupling between these regions modulates this transformation.

Importantly, however, it is important to interpret our connectivity results from PPI and DCM with caution. Prior work has described direct anatomical pathways linking RSC with both mnemonic regions (ERC, HC) as well as with sensorimotor areas^20–22,37^. However, there is limited evidence for direct GABAergic inhibitory projections from M1 to these mnemonic regions. A likely intermediary area is the posterior parietal cortex (PPC), which has extensive connections with both RSC and M1^66^. Inhibitory interactions between PPC and M1 have been suggested to relay motor-related information needed for movement control^7,67,68^ and motor imagery^69^. At the same time, PPC also projects strongly to RSC, where it may support allocentric–egocentric transformations^66^. Future studies could directly investigate the role of PPC in modulating inhibitory effects between M1 and mnemonic regions.

We find that the geometry of mnemonic force-time cognitive maps is shaped by learning and motor perceptual bias. Participants who learned the force-time space quickly across training developed less warped force-time space in mnemonic regions, whereas participants with a stronger perceptual bias towards force exhibited more warped force-time spaces. Importantly, these influences were region - specific within the mnemonic network. Warping in the HC was selectively related to learning rate, consistent with its role in stabilizing and consolidating learned relational structure^41,44–46^. In contrast, warping in the ERC was selectively associated with motor perceptual bias. Prior work shows that distortions in the environment, such as changes in geometry^47,70^, reward placement^71,72^, or barriers^19,42^, can systematically alter how ERC encodes physical space, and that the ERC provides a structure for place cell formation in HC^73–76^. Together, these findings suggest that ERC representations could be particularly sensitive to how a learned space is utilized and experienced, whereas HC representations could reflect how the space itself is constructed through learning. The RSC showed sensitivity to both learning and motor perceptual bias, highlighting its integrative role in processing both sensorimotor and mnemonic information^20–22,37^. Collectively, these results indicate that sensorimotor cognitive maps are not uniform across individuals and can be sculpted through experience.

Our results further demonstrate that mnemonic regions implement distinct computational processes when navigating sensorimotor space. The HC reflected individual differences in sensitivity to traversal distance in the learned force-time space, mirroring findings showing that hippocampal representations can be continuously updated through path integration^48–50,77^ during spatial navigation. The HC may use similar computational mechanisms to update one’s position as individuals mentally traverse a learned sensorimotor space. In contrast, the RSC exhibited a distinct computational role in integrating sensorimotor feedback with task-relevant cues. The RSC has been implicated in meditating transformations between egocentric and allocentric reference frames^20,24,78–80^. The RSC could similarly support a transition between endogenous, effort-weighted representations of force-time space and exogenously defined, task-relevant representations. Overall, these results show that mnemonic regions implement complementary computational processes. While the HC could support navigation through a learned task-relevant sensorimotor space, the RSC could integrate ongoing sensorimotor feedback with this learned space to guide communication between sensorimotor and mnemonic regions.

In conclusion, our results indicate that cognitive mapping in mnemonic regions interact flexibly with the sensorimotor system to support task-dependent movements. These mechanisms may clarify how motor and associative deficits develop in patient populations. For example, patients with mild cognitive impairment (MCI) and Alzheimer’s disease (AD), often characterized by mnemonic issues, also show motor impairments^81–84^. Additionally, AD patients display decreased activity in SMA and abnormal hyperconnectivity between M1 and cortical regions^85^, indicating disrupted communication of sensorimotor information across networks. Similarly, patients with chronic fatigue such as Myalgic Encephalomyelitis/Chronic Fatigue Syndrome (ME/CFS) report increased effort perception^86^, which could reflect impaired transmission of sensorimotor information to various cortical regions. Overall, our findings highlight the importance of the interaction between mnemonic and sensorimotor systems for understanding the emergence of motor dysfunction in clinical populations. By linking sensorimotor control with cognitive mapping, this work opens new possibilities for identifying neurological biomarkers and creating interventions that address both motor and cognitive decline.

## Methods

### Experimental setup

The game environment and behavioral data were presented and recorded using custom MATLAB scripts (http://www.mathworks.com) with the PsychToolBox library (Brainard, 1997). An MRI-compatible screen displayed the visual stimuli, which participants viewed through a 45 - degree inclined mirror positioned above their heads.

An MRI compatible force-grip sensor (TSD121B-MRI, BIOPAC Systems, Inc., Goleta, CA) was used to record hand grip isometric exertions. During experiments, signals from the sensor were sent to the custom designed software for real-time visual feedback of participants’ exertion. During the behavioral training sessions, participants performed effort exertions by holding the force-grip sensor in their right hand while sitting. During sessions in the fMRI scanner on the final day, participants held the force-grip sensor in their right hand with arm extended while in a supine position.

For recording responses during the motor and behavioral choice tasks, participants used an MRI - compatible response box (Cedrus RB-830, Cedrus Corp., San Pedro, CA) with three horizontally aligned buttons, held in their left hand.

### Experimental Procedures

#### Participants

All participants were right-handed and were prescreened to exclude those with any history of neurological or psychiatric illnesses. This study was approved by the Institutional Review Board at the Johns Hopkins School of Medicine, and all participants provided informed consent.

Forty participants were enrolled in the experiment. Seven participants discontinued participation in the study due to various reasons : three participants found the force grip exertions too painful and had to stop the study during day 1, four participants had difficulty understanding the instructions and had to stop the study on day 1. The final analysis included N = 33 participants in total (mean age, 23.7 years; age range, 19-34 years; 19 females).

#### Force-Time Space

Throughout the experiment, participants were exposed to exertions with different force and time levels. The four force levels of exertions were arranged linearly from 10 to 55 % of participants’ maximum voluntary contraction (MVC). The time levels of exertions were arranged linearly from 1.5 to 5.5 seconds (1.5, 2.83, 4.17, 5.5 s). Each cue (Cues 1-16) had a unique combination of force and time levels and was arranged in a 4 × 4 grid-like force-time space.

#### Cue presentation

The presented cues consisted of 16 grayscale images of fruits and vegetables. These were obtained free to use for digital or printed media on the https://www.flaticon.com website. To ensure that the visual features of stimuli (e.g., outline thickness, curvature) were not associated with the exertion information, the pairings of each cue and their respective force and time levels were randomly assigned to each participant.

#### Visual feedback of exertions

During exertion trials, participants were instructed to perform an isometric exertion and guide a cursor to a central target. The cursor moved vertically proportionally to the amount of force exerted. Upon reaching the target, the cue next to the central target begun filling up in color. During this time, participants had to maintain the exertion. Once the cue was filled, participants had to release their grip for the cursor to return to baseline. Greater force levels required higher exertions to get to the central target whereas greater time levels required participants to hold the exertion for a longer time. For a successful exertion trial, participants had to fulfill three requirements. First, participants needed to reach the target within four seconds. Second, during exertion, the cursor needed to stay within the central target (+/- 3 E.U.) for more than 50% of the time. Finally, the cursor needed to return to baseline within +/- 0.5 seconds of the required time level. If the requirements for a successful exertion were met, participants were provided with feedback (“Success”). If not, feedback was provided (“Failure”) and participants had to redo the exertion until success.

#### Experimental paradigm

The experimental paradigm consisted of a multi-day study (**Supplementary Fig. 1**) where participants alternated between 1) behavioral training sessions to associate isometric hand-grip exertions with external cues and 2) assessment sessions that evaluated their ability to integrate learned cue-exertion pairings with sensorimotor feedback. The methodological design for the sessions during each experimental day is explained in detail below.

### Day 1

#### Behavioral Training Session

During behavioral training, participants alternated between performing isometric exertions on the force-grip sensor and the assessment phase (**Supplementary Fig. 3**). Participants began each behavioral training session by performing an isometric exertion to guide a cursor to a central target. During this process, participants were instructed to associate the force or time level of the exertion with the cue that appeared next to the central target. For ease, participants were instructed to focus on one dimension (force/time). Of the final participant cohort (n=24), half the participants focused on learning the force levels during day 1 behavioral training while the other half focused on learning the time levels. Following a series of exertions, participants were then presented with the assessment phase where they were presented with two cues they experienced during exertion and had to select the cue that was greater in either the force or time level (dependent on the dimension they were getting trained on). The behavioral training session was divided into three chunks : chunk 1, chunk 2, chunk 3. During Chunk 1, participants completed four sessions where within each session, they were exposed to four cues, where each cue was repeated three times, resulting in a total of 12 exertion trials. Cues were such that they would differ in the dimension the participants were instructed to focus on but be constant in the other dimension (**Supplementary Fig. 3B**). Cues were presented in a pseudorandom order from three possible sequences. For example, during time training on cues 1-4, the possible sequences would be: 1-2-4-3, 3-4-2-1, 2-3-4-1. Following the 12 exertion trials, participants completed the assessment section where we sampled pairwise adjacent cues within the four cues. Each pair was asked twice, resulting in a total of 6 questions per session. Chunk 2 followed the same format as Chunk 1, with the difference being that participants were exposed to 8 cues at once, resulting in two sessions. During the assessment, pairs were now sampled from adjacent, diagonal pairs of cues within the session. Finally, on Chunk 3, participants were exposed to random sequence of cues and during assessment, they were tested on diagonal cue pairs that were adjacent in the tested dimension and not asked during assessment phase in Chunk 2 (**Supplementary Fig. 3B**).

#### Motor Space Navigation Task

During the motor space navigation (MSN) task, participants completed two segments: 1) localization and 2) navigation. During localization, participants performed an isometric exertion but this time, the cue identity was hidden by replacing the cue next to the central target with an arbitrary circle. During the exertion process, participants were instructed to identify which of the 16 cues the circle best represented based on the experienced force and time level. Following the exertion, participants were exposed to the navigation segment, where they passively viewed a target cue that was different from the hidden cue. During this navigation segment, participants were instructed to compute the difference in force and time levels between the hidden cue that they had to identify during the localization, and the target cue. This was equivalent to traversing and computing the distance between the two cues in the force-time space. This process was then repeated a second time with the same hidden cue during localization, but a different target cue in the navigation segment. Finally, participants selected the target cue that was further from the hidden cue in the force-time space, which required comparing the two trajectories. The sampling of cues for the localization and navigation segments was adapted from Park et al., 2021, which utilized a similar design for testing cognitive map organization of social hierarchies^26^. A session consisted of 48 trials, which resulted in participants having to compute 96 trajectories to perform the MSN task. Each segment was separated by a fixation cross.

Participants started day 1 by performing a maximum voluntary contraction (MVC), which allowed us to scale the force levels depending on their maximum grip strength. Following MVC, participants performed the first MSN session, which enabled us to get a baseline measure of their performance without having been exposed to the force-time space. They then completed the behavioral training session on one dimension and ended the day by performing the second MSN session and MVC.

### Day 2

Day 2 largely followed the same format as day 1 except during the behavioral training session, participants were instructed to focus on the second dimension (i.e., participants that completed the force dimension training on day 1 would do time training). Following the second MSN session on day 2 and MVC, participants completed the subjective decision-making and spatial reconstruction task, which are described in detail below.

#### Subjective Decision-Making Task

Participants were shown two cues and had to select the cue that felt less effortful to exert after considering differences in the force and time levels between the two cues. Participants completed a total of 120 trials which comprised of all possible combinations of cue pairs. At the end of the experiment, participants were informed that ten of the trials would be selected at random and participants would have to successfully perform the exertions of the cues that they had selected in the selected trials.

#### Spatial Reconstruction Task

Participants were presented with all 16 cues that were organized in randomized locations on the left side of the screen. The right side of the screen consisted of an empty force-time grid with the x-axis representing the force level and the y-axis representing the time level. Participants were instructed to reconstruct the learned force-time space from memory by dragging each cue into their respective locations. Prior to the spatial reconstruction task, participants were never exposed to the full force-time space.

### Day 3

Day 3 largely followed the same format as day 2 except during the behavioral training session, participants were instructed to review both dimensions. Specifically, during chunk 1, participants did a refresher of the first dimension that they learned on day 1 that mirrored chunk 2 of day 1 (**Supplementary Fig. 3C**). During chunk 2, participants did a refresher for the second dimension that mirrored chunk 2 of day 2 training. For both chunks, each cue was only shown once, resulting in a total of 16 trials. In chunk 3, participants were exposed to all 16 cues presented in random order. During the assessment phase, participants were asked to report which of the presented cues had a greater force or time level depending on the prompt. Each pair was thus presented twice (one asking for force level, one asking for time level). The cue pairs for chunk 3 were selected as adjacent diagonal pairs in the force - time space (**Supplementary Fig. 3C**). In addition, following the second MSN session on day 3, participants completed the localization task (described below). They then ended the day by performing the subjective decision-making and spatial reconstruction task. Participants’ performance in the second MSN session for day 3 had to be greater than 80% success rate for them to be brought back for the sessions in the MRI scanner on day 4.

#### Localization-only Task

Participants performed an isometric exertion that mirrored the localization segment of the MSN task, where the cue was replaced with an arbitrary circle. Resembling the MSN task, participants had to identify which of the 16 cues best matched the exertion. Following exertion, participants were presented with a screen that showed the correct cue as well as seven other distractor cues. Participants had to select the cue they thought best matched the exertion profile. If the correct cue was selected, participants were provided feedback (“Correct”), and the selected cue would no longer show up as the correct cue but still had the possibility of showing up as a distractor. If the incorrect cue was selected, participants were provided feedback (“Incorrect”), and the trial would be sent to the end of trial presentation, which meant that participants would have to eventually identify the correct cue again. Participants had to correctly identify all 16 cues to finish the localization-only task.

### Day 4

On Day 4, participants initially performed an MVC and completed a behavioral training session with the goal of getting accustomed to performing isometric exertions while lying inside the MRI. The training structure was identical to the one in day 3, with the exception that participants did not perform the assessment segments. Following training, participants performed three sessions of the MSN task while being scanned in the MRI machine. Each session was split into three blocks, which resulted in participants completing a total of 9 blocks. Following the end of the MSN sessions, participants provided a second MVC to end the experiment.

#### MRI protocol

The MRI scanning sessions were conducted on a 3 Tesla Philips Achieva Quasar X-series MRI scanner equipped with a radio frequency coil. High-resolution structural images were acquired using a standard MPRAGE pulse sequence, covering the entire brain at a resolution of 1 mm × 1 mm × 1 mm. Functional images were collected at a 30° angle from the anterior commissure-posterior commissure (AC-PC) axis, with 48 slices acquired at a resolution of 1.87 mm × 1.87 mm × 2 mm to ensure full brain coverage. An echo-planar imaging (FE EPI) pulse sequence was applied (TR = 2800 ms, TE = 30 ms, FOV = 240, flip angle = 70°).

#### Pre-processing

SPM12 software package was used to analyze the fMRI Data (Wellcome Trust Centre for Neuroimaging, UCL, London, UK). A slice-timing correction was applied to the functional images to adjust for the fact that different slices within each image were acquired at slightly different time points. Images were corrected for participant motion by registering all images to the first image, spatially transformed to match a standard echo-planar imaging template brain to account for anatomical differences between participants. Finally, the images were smoothed using a 3D Gaussian kernel (8 mm FWHM). For certain analyses (e.g., testing for hexadirectional modulation in the ERC, Pattern Component Modeling), non-smoothed images were used to preserve fine-grained voxel-level differences and to account for small regions of interest (e.g., ERC). The final resolution of the preprocessed images had voxel dimensions of 2.00 mm x 2.00mm x 2.00mm. This set of data was then analyzed statistically.

### Behavioral Analyses

#### Modeling spatial metrics during navigation of force-time space

Prior to the conducting the analysis, trials in which the reaction time deviated more than 2 standard deviations from the mean were removed (∼5% of trials). In addition, the analysis only included MSN trials after the training session on day 1, as on the first day before training, participants were not exposed to the force-time space. Participants’ response in the MSN task was modeled using a mixed-effects logistic regression in the following form:

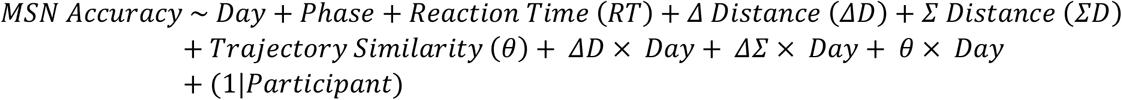

Where MSN Accuracy is a binary variable that represents trial-by-trial accuracy (Correct = 1, Incorrect = 0). Δ Distance was computed by taking the absolute difference in Euclidean distance between two trajectories traversed in the MSN trial:

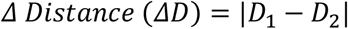

Σ Distance is computed by taking the sum in Euclidean distance between the two trajectories in the MSN trial:

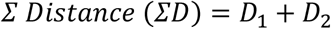

Finally, Trajectory Similarity was computed by taking the angle (*θ*, [0, 180 °]) between the two trajectories. In addition to these three spatial metrics (ΔD, ΣD, *θ*), the model also contained experimental day (Day = 1,2,3,4), phase number (Before training = 1, After training = 2), reaction time (RT), and the interaction between spatial metrics and experimental day. Participant ID was entered as random effects. All metrics were z-scored.

To determine how participants weighted differences in force and time levels when performing the MSN task, a logistic regression was used to predict performance in the following form:

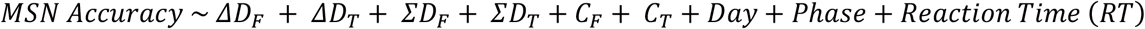

Where each spatial metric is now computed separately for the force and time axes instead of taking the Euclidean distance (See **Fig. 4A, B**). Trajectory similarity for the force and time dimensions as computed by taking the distance between navigating cues in the force (*C*_*F*_) and time (*C*_*T*_) dimensions. This process was repeated for each participant, and the coefficient estimates for each spatial metric and the Wilcoxon paired signed-rank test was used to test if across the group level, if there was a difference between the spatial metrics in the force vs. time dimensions.

#### Modeling motor perceptual bias in subjective decision-making task

Each participants’ in the subjective decision-making task for days 2 and 3 were combined and modeled using logistic regression in the following form:

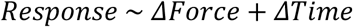

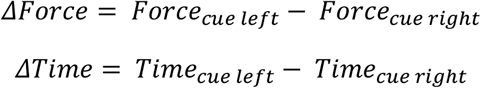

Where response is a binary variable that represents whether the participant selected the left cue (1) or right cue (0). Δ Force was computed by taking the difference between the force levels of the left and right cues. Δ Time was computed by taking the difference between the time levels of the left and right cues. Regression coefficients for Δ Force and Δ Time were computed for each participant and a paired t-test was used to test for significant differences between the coefficients. Motor perceptual bias was computed for each participant by taking the difference between Δ Force and Δ Time.

#### Modeling learning rate

Each participants’ accuracy in the MSN task across individual sessions was modeled using an exponential function in the following form:

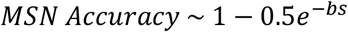

Where s represents session number and b reflected the learning rate. The fitnlm function in MATLAB was used to fit the exponential function for each participant. Participants’ performance in the first session of the MSN task was excluded as participants had not been exposed to the force-time space and made random choices during the session.

### fMRI Analysis

#### Contrasting brain activity during localization and navigation

A general linear model (GLM) was used to generate voxel-wise statistical parametric maps from the fMRI data. A GLM was specified that included conditions spanning different segments of the MSN task. The outline of the GLM that was constructed for each participant is shown below:

### GLM 1

1. **Localization segment** : exertion (∼2-8 s)
2. Exertion feedback (1 s)
3. **Navigation segment** : cue presentation trials (4-6 s)
4. Presentation of the decision screen (∼2 s)
5. Time of button press during decision screen (modeled as event regressor)

We defined the duration of the localization segment to cover the entire exertion profile, starting from the time at which the trial started until the participant released the exertion. Exertion feedback was modeled separately as an event regressor. The presentation of cues during the navigation segment was modeled as a block regressor where the onset was set as the initial presentation of the cue and the duration depended on the total time the cue was presented to participants (4-6 s). The decision segment was modeled using two separate regressor. The first regressor included the onset when the decision screen was first presented. The second regressor indicated the time of button press. GLM 1 enabled us to contrast whole-brain activity at the time of localization and navigation.

#### Region of interest (ROI) specification

ROIs for the hippocampus (HC) and entorhinal cortex (ERC) were obtained from Park et al., 2021^26^. The ROI for the retrosplenial cortex (RSC) was constructed by combining Broadman areas 29 and 30, which were obtained from the WFU PickAtlas^87,88^ and dilating the mask by a factor of 1.5 to ensure adequate coverage. The ROIs for the primary motor cortex (M1), cerebellum (CBL) and supplementary motor area (SMA) were obtained by taking a conjunction between an anatomical mask for each region, which were obtained from the WFU PickAtlas^87,88^, and a group-level Localization – Navigation contrast set at p < 0.001 (**Supplementary Fig. 7A**). For running ROI analysis on contrasting activity of sensorimotor brain areas during localization and navigation as well as functional connectivity, we utilized the leave-one-subject-out (LOSO)^89^ approach to prevent bias since the sensorimotor ROIs were obtained from the same whole-brain contrast. The LOSO approach mirrored the same conjunction, with the exception that the contrast was obtained from all but one participant. The overlap between the functional and anatomical mask would then be used as the sensorimotor ROIs for the left-out participant.

#### Testing for hexadirectional modulation in the ERC

We leveraged established fMRI methods for testing grid-like activity in the ERC as participants navigated the force-time space^26,29,34^. The 6-fold signals in ERC population activity are represented as: cos(6(*θ* − *ϕ*)), which can then be decomposed into: cos(6*θ*) ∗ cos(*ϕ*) + sin(6*θ*) ∗ sin(*ϕ*). *θ* is the direction of travel from the hidden to target cues in the force-duration space and *ϕ* is the grid orientation. To estimate *ϕ*, we constructed a second GLM (GLM 2), which had the same main regressors as GLM 1 but additionally assigned cos(6*θ*) and sin(6*θ*) as parametric modulators at time of cue presentation. We then extracted *β*_*cosine*_ and *β*_*sine*_ estimates for the ERC, which would correspond to estimated values of cos(6*ϕ*) and sine(6*ϕ*). *ϕ* can be estimated using the function: arctan 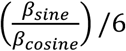. Rayleigh’s test was used to test for non-uniform distribution of putative grid orientations estimated from each fMRI block across voxels for every participant. To test for 6-fold activity, we constructed GLM 2, which assign ed a parametric modulator: cos(6(*θ* − *ϕ*)) at time of cue presentation using *ϕ* estimated from GLM 1. From GLM 2, we obtained participant-specific *β*_*grid*_, corresponding to the estimate for cos(6(*θ* − *ϕ*)). For this estimation process, we implemented a 9-fold cross-validation (CV) procedure by splitting the fMRI sessions into a training and testing dataset. Since there were a total of nine fMRI blocks, the training data would consist of 8 blocks and the testing data would consist of the left-out block. The training data was used to estimate *ϕ* from GLM1 and implemented on the testing set in GLM 2. This process was repeated 9 times where each block served as a test set once. To test grid-like signals in the ERC, we utilized a one-sample Wilcoxon signed-rank test whether *β*_*grid*_ estimates area significantly different from 0 due to the non-normal distribution of *β*_*grid*_. In addition, we repeated the analysis described above with models assuming different control periodicities (n = 4,5,7,8), which resulted in*β*_*grid*_ that describe how much the ERC activity was explained by these control periodicities. We then utilized paired Wilcoxon signed-rank test to determine whether the 6-fold periodicity signals in ERC were significantly greater than that of control periodicities.

To better visualize the 6-fold periodicity signal in ERC, we investigated whether ERC activity would be greater for trajectories that were aligned to *ϕ*. Using the cross-validated *ϕ* estimates from GLM 1, we binned the corrected trajectories (*θ* − *ϕ*) into 12 equal bins of 30 °. We then created GLM 3, which modeled each trajectory bin as a separate regressor. For every participant, we then extracted and visualized ERC activity for each trajectory bin. We utilized a paired t-test to determine whether the mean ERC activity for aligned trajectories was significantly greater than mean ERC activity for misaligned trajectories. For visualizing the force-time grid code, we first took the average ERC activity modulo 180 ° (e.g., averaging ERC activity for 0° and 180°) due to the isotropic nature of grid fields. The averaged values were then interpolated into polar coordinates.

#### Pattern component modeling (PCM)

We implemented PCM^36^ to investigate whether the multi-voxel activity patterns within our priori sensorimotor and mnemonic ROIs are explained by the force-time space. Specific details on implementing PCM can be found in Diedrichsen et al., 2018^36^. PCM models brain activity data Y, which is a matrix consisting of N time points (trials) and P voxels, using the following equation:

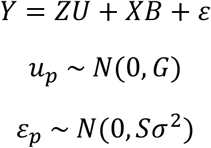

Where U is the matrix of true activity patterns (number of conditions x number of voxels) and Z is a design matrix (trial number x number of conditions). PCM also models effects of no interest (B), which includes the condition-independent mean activity pattern, as well as noise (ε). The activity patterns (u_p_) are random variables drawn from normal distribution. The noise of each voxel is modeled as Gaussian with constant term σ^2^, with the assumption that the noise is independent within each imaging run S = I. The goal of PCM is to construct a representational model that describes the covariance matrix of the activity profiles (G). We implemented component models where the covariance matrix was modeled as a linear sum of weighted component matrices:

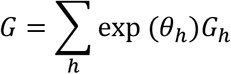

We constructed predicted covariances that described the representational structure of each cue due to differences in force (force component matrix) and time (time component matrix) levels. Each component matrix was computed by taking the inner product of the feature set *FF*^*T*^ where F corresponded to either the force or time levels (ranging from level 1 -4) of the cues. We leveraged the eight most frequently sampled cues during the localization segment (**Supplementary Fig. 4A, Fig. 3D**, yellow). We specified three different component models: A force only model (F), which modeled the neural covariance structure based on the force component, a time only model (T), which modeled the structure based on the time component, and a force + time model (F+T), which modeled the structure using a weighted linear combination of both force and time component matrices. In addition to the component models, a null model was specified which set G = 0, which assumed no difference between activity patterns across different cues. Finally, a noise ceiling (free) model was specified which predicted G without any constraints. The unconstrained G was modeled as:

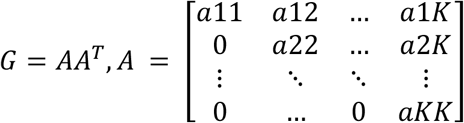

Where A is an upper-triangular matrix where free parameters correspond to non-zero elements of A. In our case, since we had 8 total cues included for the analysis, we assigned K = 8, which resulted in a total of 36 parameters. Maximum likelihood estimation through conjugate gradient descent was used to optimize model fit to the data. To evaluate model performance, we implemented cross-validation across participants where model parameters are fitted to n-1 subjects, using separate noise and scale parameters to each subject, and subsequently evaluated on the left-out subject. The cross-validated log-likelihood values of the three components were scaled with respect to the null model. The performance of the noise ceiling model to the group data indicated the upper limit of model performance given the noise structure within a region while its performance in the cross-validated setting provided a lower bound to the noise ceiling. Parameter estimates for the F + T model, which corresponds to how much weighting was assigned to the force (ω_F_) and time (ω_T_) covariance matrices when estimating the neural covariance structure, were obtained by fitting the model individually to each participant individually. For visual purposes, the predicted and neural covariance matrices were centered by assigning the mean of rows and columns to zero, which was equivalent to subtracting out the mean pattern.

#### Psychophysiological interaction analysis (PPI)

A PPI analysis was performed to investigate the functional coupling between sensorimotor and mnemonic regions during the MSN task. We assigned the seed region as the RSC to investigate the a priori hypothesis that the region acts as an integrative region facilitating transformation of force-time information between sensorimotor and mnemonic brain regions. The PPI analysis was performed by adapting GLM 1 to include two additional regressors:

1. A regressor that contains a representative time course of activity in RSC (seed)
2. Interaction between RSC activity and the Navigation – Localization contrast.

The interaction regressor allowed us to investigate whether any other brain region showed significantly greater functional coupling with RSC in the Navigation vs. Localization conditions. ROI analysis was performed for the mnemonic and sensorimotor regions to identify significant coupling with RSC.

#### Dynamic causal modeling (DCM)

We implemented a deterministic, bilinear DCM to test directed causal interactions between the sensorimotor and mnemonic brain regions. DCM utilizes forward modeling to represent how experimental manipulations lead to change in neural activity through a bilinear differential equation in the form of:

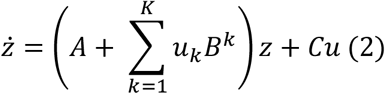

where ż represents the change in neural activity per unit time (i.e., derivative of neural activity for each region), u represents experimental conditions (i.e., successful or failed task performance outcomes), A is a matrix that defines the intrinsic coupling (average effective connectivity) between regions, B is a matrix specifying the modulation of effective connectivity due to experimental condition k = 1…K, and C is a matrix encoding the sensitivity of each region to driving input from experimental condition k = 1…K. Each connection thus represents a change in neural activity due to directed signals (A, C matrix), or the modulation of the directed connection (B matrix).

We followed the procedures described in Zeidman et al. for specifying DCM across participants^38,39^. Given what we observe from Navigation – Localization contrast and PPI results, we assigned input nodes (seed regions) as RSC, M1, SMA and HC. We enabled RSC and HC to be driven by the Navigation condition while M1 and SMA were driven by the Localization condition (i.e., C matrix). We enabled full bidirectional connections between all regions (i.e., A matrix). For the DCM analysis, we were specifically interested in parameters modulating the modulation between all regions following the Localization and Navigation conditions (i.e., B matrix).

#### Parametric Empirical Bayes (PEB)

The PEB approach uses hierarchical Bayesian modeling where each parameter (i.e., the estimate of connection strength) is treated as a random effect. All participants are assumed to have the same model architecture but varying strengths of connections within the group^39^. Thus, one full DCM is specified per participant, enabling all connections, and a series of candidate models to be specified that test different connectivity hypotheses. Bayesian Model Reduction (BMR) was applied to identify a reduced model that provides the most parsimonious explanation of the observed data. To do so, model evidence (Free energy) is computed which measures the trade-off between the accuracy (i.e., ability to predict observed BOLD activity) and complexity (KL-divergence between estimated parameter and the original priors)^38,90^. BMR iteratively discarded connection parameters that did not contribute to model evidence until the point at which discarding any parameter began to decrease model evidence. A Bayesian Model Average (BMA) was then calculated over the 256 models from the final iteration of BMR, where the average of the parameters from different models are weighted by the models’ posterior probabilities^91,92^. We present and interpret the results of PEB parameters with very strong (p > 0.99) Bayesian evidence^93^.

For our first DCM analysis, the PEB design matrix included just a constant term, which represented the mean coupling strength between sensorimotor and mnemonic brain regions following the navigation and localization conditions. For the second DCM analysis, the PEB matrix included both a constant term and mean-centered downregulation of force in the RSC:

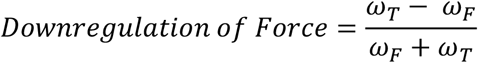

Which we computed for each participant using the ω_F_ and ω_T_ values obtained from the PCM analysis. Since we were specifically interested in whether the connectivity parameter from M1 to RSC during localization was significantly modulated by the time weighting index in RSC, we computed the posterior probability of the connection parameter being greater than 0. Following PEB estimation, we utilized leave-one-out (LOO) cross-validation using the observed effects of interest (i.e., modulation from M1 to RSC during localization) to estimate the out-of-sample downregulation of force in RSC for each participant. The LOO approach provides a statistically robust approach to assess the validity of associations^39^. We report Kendall’s rank correlation (τ) and the p-value for the right-tailed correlation between the model estimated and observed downregulation of force in RSC. We then repeated the second DCM but this time, investigated whether the M1 to HC connectivity was modulated by the downregulation of force in HC.

#### Mediation analysis to test association between motor-perceptual bias, M1-Mnemonic Connectivity, and mnemonic force-time representation

For each participant, M1-Mnemonic connectivity was computed by taking the mean of M1-RSC and M1-HC connectivity parameters obtained from the first DCM analysis. In addition, mnemonic force downregulation was computed by taking the mean force downregulation in RSC and HC for each participant. We then performed a mediation analysis to investigate the relationship between motor perceptual bais (X), M1-Mnemonic connectivity (M), and force downregulation in mnemonic regions (Y). Mediation analysis was conducted using the Mediation Toolbox (Matlab code for mediation analyses is freely available at: https://github.com/canlab/MediationToolbox). Given the non-normal distribution of the variables, we employed robust regression with bootstrapping (10,000 resamples), which allowed us to assess direct and indirect effects using percentile bootstrap confidence intervals using significance threshold set at p < 0.05.

#### Associating neural representation of force-time space with learning rate

We first quantified a warping metric for each participant that defined the degree to which the force-time space was warped:

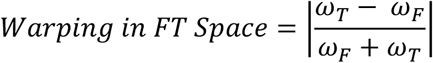

Where ω_F_ and ω_T_ are the weighting of the force and time covariance matrices obtained from the PCM analysis. Values closer to 0 indicated an unwarped representation of force and time whereas values closer to 1 indicated a more warped representation. To determine whether participants’ learning rate or motor perceptual bias influenced the warping of mnemonic force-time space, we utilized a robust linear regression in the following form:

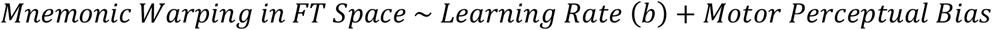

Where mnemonic warping in FT space was computed by taking the average warping in force time space for the RSC, HC and ERC. To plot and visualize the effects for learning rate, an adjusted response function was utilized that displayed the relationship between mnemonic warping and learning rate while controlling for the effects of motor perceptual bias. Similarly, an adjusted response function was used to visualize the effect of motor perceptual bias on mnemonic warping while controlling for the effects of learning rate. The same regression framework was utilized to model the warping in FT space separately within sensorimotor regions, and within each mnemonic ROI.

#### Associating neural representation of force-time space with behavioral measures when navigating sensorimotor space

We computed the bias for each spatial metric computed earlier in the behavioral analysis of MSN performance, defined as the difference between parameters derived from the force and time dimensions:

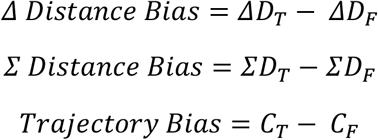

We first computed Kendall’s rank correlation to find preliminary evidence of potential association between the spatial bias metrics and downregulation of force in mnemonic force-time space. We found that after correcting for multiple comparisons (n = 9 correlations, Holm-Bonferroni method), Σ Distance Bias was significantly correlated with downregulation of force in the HC. We then conducted a robust linear regression in the following form:

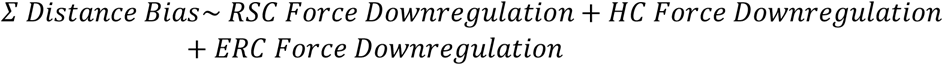

An adjusted response function was used to visualize the effect of HC Force Downregulation on Σ Distance Bias while averaging out the effects of RSC and ERC Force Downregulation.

To test whether force-time space in RSC reflected sensorimotor and task-related processing, we utilized a robust linear regression to model participants’ performance in the localization-only task in the following form:

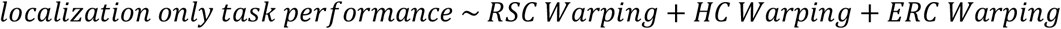

Where localization-only task performance, defined as each participants’ accuracy in the task, was modeled as a function of their warping metric for the RSC, HC and ERC. An adjusted response function was used to visualize the effect of RSC warping on participants’ performance in the Localization-Only Task while controlling for the effects of warping in HC and ERC.

To test whether the errors in localization-only task were related to how force and time were represented in the RSC, force downregulation in RSC was modeled as a function of errors made along each dimension during the localization-only task using a robust linear regression in the following form:

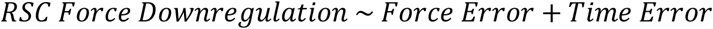

Where Force and Time errors were computed by taking the average distance between the correct cue and localized cue in the force and time dimensions respectively across all localization-only trials. An adjusted response function was used to visualize the effect of force errors on force downregulation in RSC while controlling for the effect of time errors.

