## Supplementary Information for "Transformations of cognitive maps for sensorimotor control"

[1] Department of Biomedical Engineering, Johns Hopkins School of Medicine, Baltimore, MD 21205, USA. [2] Kennedy Krieger Institute, 707 North Broadway, Baltimore, MD 21205, USA. [3] Department of Neuroscience, Johns Hopkins School of Medicine, Baltimore, MD 21205, USA. [4] Neuroscience Programme, Champalimaud Research, Lisbon 1400-038, Portugal. [5] Kavli Neuroscience Discovery Institute, Johns Hopkins University, Baltimore, MD 21205, USA.

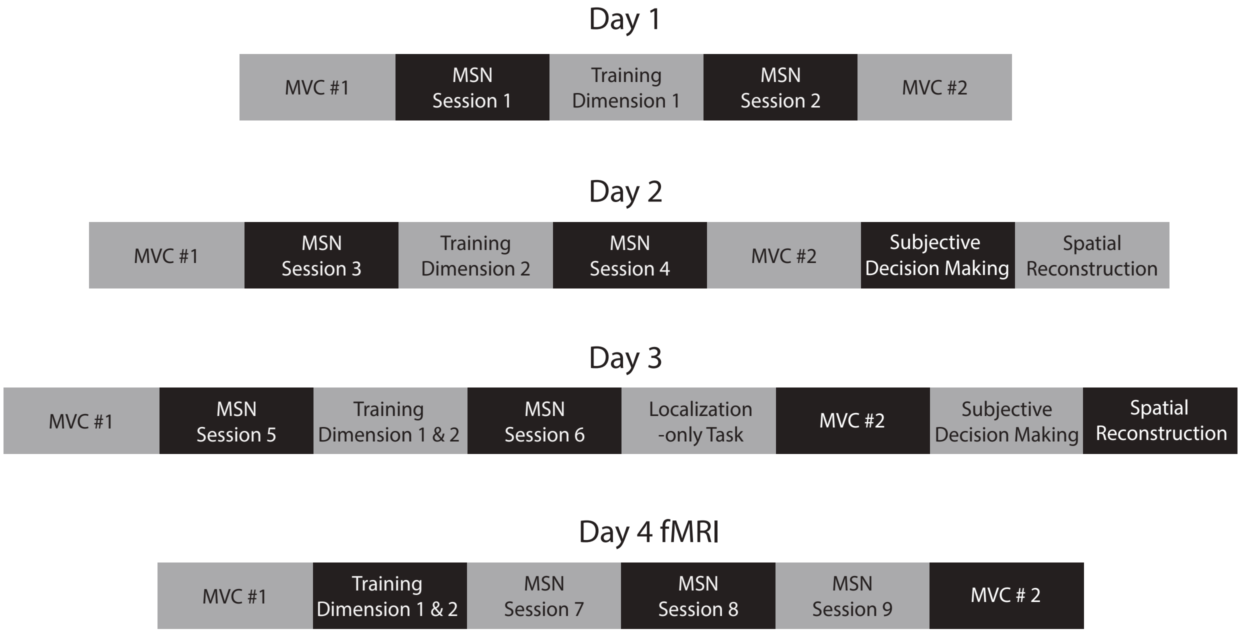


**Supplementary Figure 1: Experimental schedule**

Participants underwent a multi-day experimental paradigm. At the start of each day, participants performed a maximum voluntary contraction (MVC) which enabled us to normalize force levels based on their grip strength. Days 1-3 were performed outside the fMRI scanner and was designed to train and assess participants' ability to associate exertions with the cues in the force-time space. On Day 1, participants performed the MSN task before and after the training session. During training, participants focused on one dimension at a time where half the participants (n=12) completed force dimension training while the other half of participants (n=12) completed time dimension training. Day 2 largely followed the same format as day 1 except during training, participants completed the second dimension training (i.e., participants that did force dimension training on day 1 would do time dimension training on day 2). In addition, participants also completed the subjective decision making task (**Fig. 3A**) and the spatial reconstruction task (**Supplementary Fig. 6**). Day 3 mirrored days 1 and 2, with the exception that during the training session, participants focused on reviewing both force and time dimensions. In addition, following MSN session 6, participants completed the localization-only task (**Supplementary Fig 5**). On Day 4, participants reviewed the force and time dimensions before completing three sessions of the MSN task (sessions 7-9) inside the fMRI scanner.

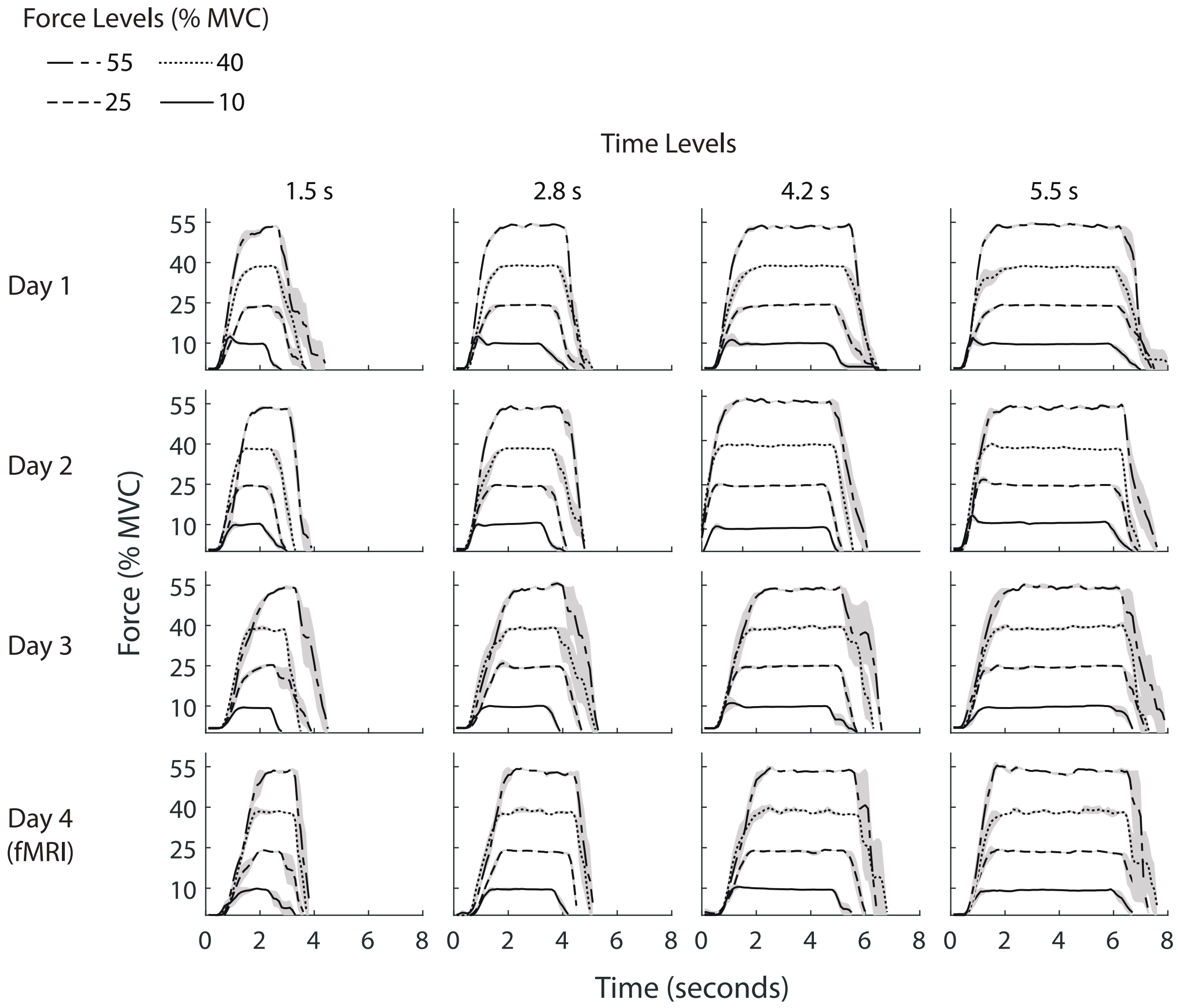


**Supplementary Figure 2: Force plots for a representative participant across training days**

Representative force trace plots for an exemplary participant across days (row) for different time (column) and force (see legend) levels. Shaded areas represent SEM.

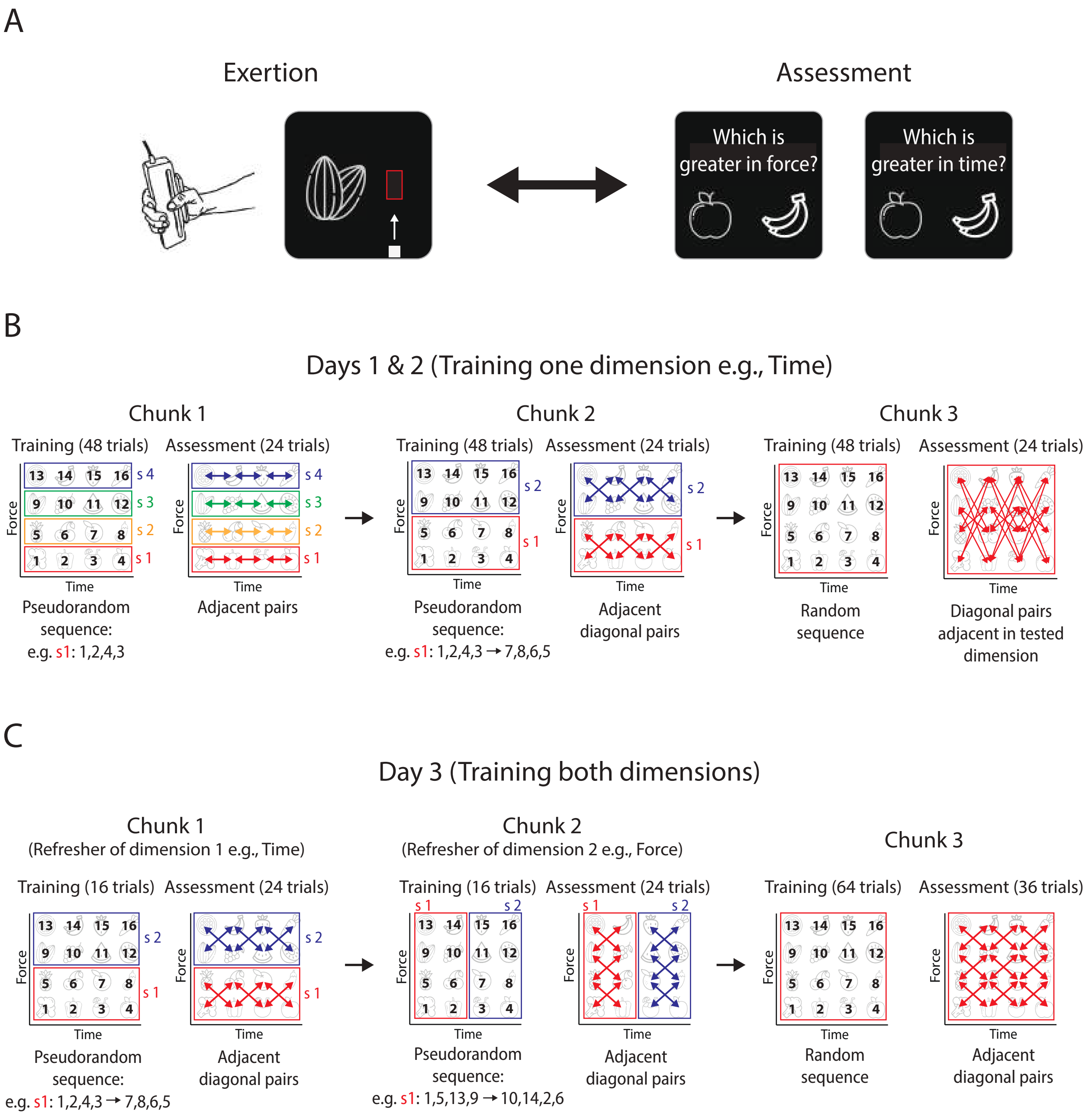
A
Exertion
Assessment
Which is greater in force?
Which is greater in time?
B
Days 1 & 2 (Training one dimension e.g., Time)
Chunk 1
Training (48 trials)
Assessment (24 trials)
Force
Time
Pseudorandom sequence: e.g. s1: 1,2,4,3
Adjacent pairs
Chunk 2
Training (48 trials)
Assessment (24 trials)
Pseudorandom sequence: e.g. s1: 1,2,4,3 → 7,8,6,5
Adjacent diagonal pairs
Chunk 3
Training (48 trials)
Assessment (24 trials)
Random sequence
Diagonal pairs adjacent in tested dimension
C
Day 3 (Training both dimensions)
Chunk 1
(Refresher of dimension 1 e.g., Time)
Training (16 trials)
Assessment (24 trials)
Pseudorandom sequence: e.g. s1: 1,2,4,3 → 7,8,6,5
Adjacent diagonal pairs
Chunk 2
(Refresher of dimension 2 e.g., Force)
Training (16 trials)
Assessment (24 trials)
Pseudorandom sequence: e.g. s1: 1,5,13,9 → 10,14,2,6
Adjacent diagonal pairs
Chunk 3
Training (64 trials)
Assessment (36 trials)
Random sequence
Adjacent diagonal pairs

**Supplementary Figure 3: Experimental protocol for training sessions**

**(A)** During training sessions, participants alternated between performing isometric exertions on the force sensor and the assessment phase. During the exertions, participants were instructed to associate the force or time level of the exertion (depending on the training day) with the presented cue. During assessment, participants were shown two cues they experienced during exertion and asked to select which cue is greater in either the force or time level (depending on training day).

**(B)** Training schedule for Days 1 & 2 followed the format where participants were instructed to focus on one dimension. For both days, the training was divided into three chunks. The visualization shown above corresponds to a training session focused on learning the time levels. For Chunk 1, participants were presented with four cues at a time for a session (e.g., s1) which varied depending on the dimension of the training day (e.g., time). Within this session, the cues were presented in pseudorandom sequence (e.g., cues 1, 2, 4, 3 for s1) three times, which resulted in a total of 12 trials per session. Following these 12 trials, participants completed the assessment section for which we sampled pairwise adjacent cues within the four cues within the session. Each pair was asked twice, which resulted in a total of 6 questions per session. Once participants finished the exertion and assessment phases assocaited with one session, they would move on to the next session. To minimizie prolonged fatigue, participants were exposed to an alternating session order (i.e., s1, s4, s2, s3) such that they would not need to exert high force or time levels for prolonged period of time. Chunk 2 followed a similar format as Chunk 1, with the difference being that participants were now exposed to 8 cues at once during training and assessment, which resulted in two sessions within the Chunk (s1, s2). During assessment, pairs were now sampled from adjacent, diagonal pairs of cues within the session. Finally, on Chunk 3, participants were exposed to random sequence of cues and during assessment, they were tested on diagonal pairs of cues that were adjacent in the tested dimension and not asked during the assessment phase in Chunk 2.

**(C)** On Day 3, participants were instructed to review both force and time dimensions during training. Chunks 1 and 2 resembled Chunk 2 that was completed on Days 1 and 2, which allowed participants to refresh their associations between the cues and their force and time levels. On Chunk 3, participants were exposed to random sequence of cues and during the assessment, were assessed on both force and time levels of the presented pairs of cues.

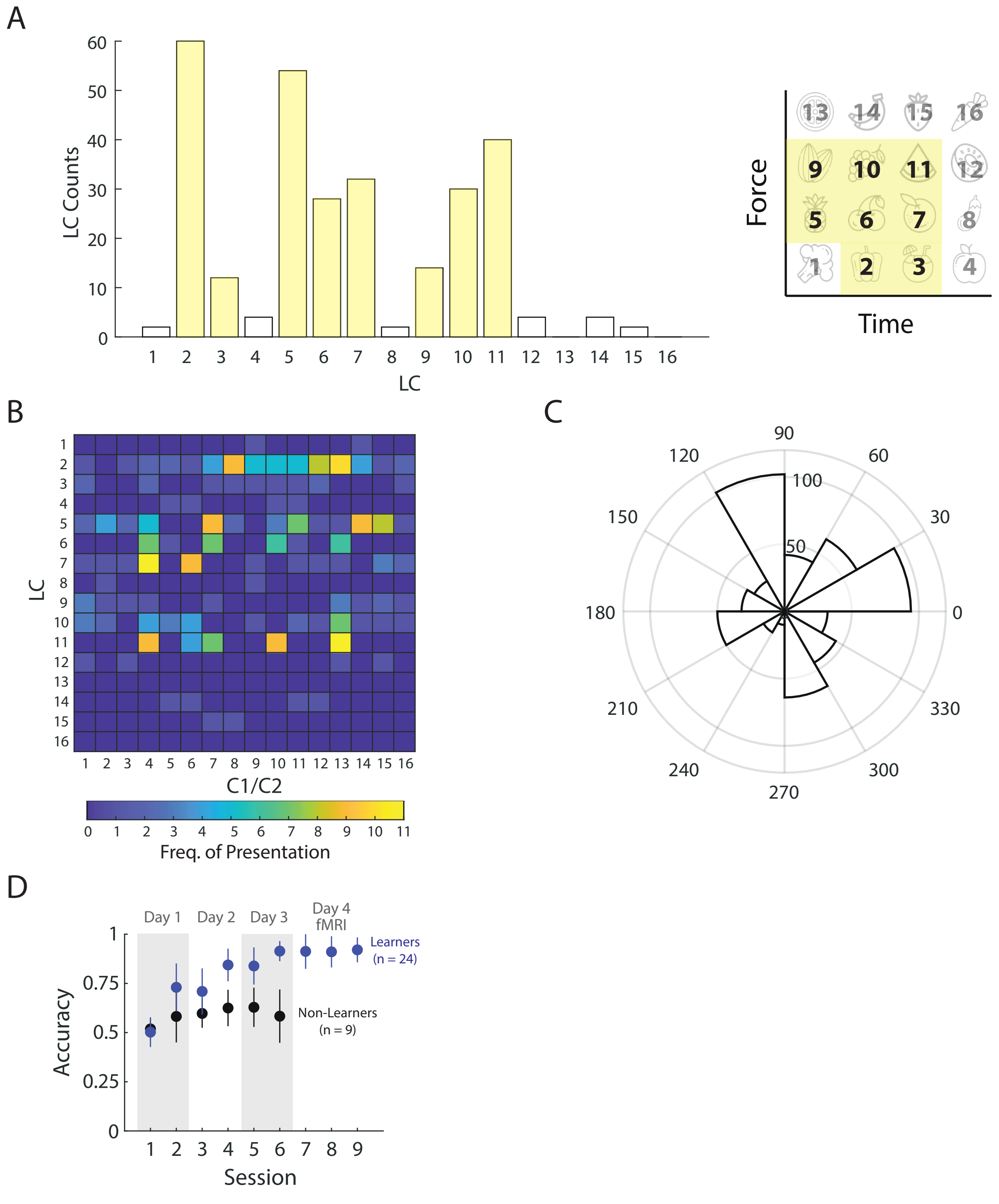


**Supplementary Figure 4: Distribution of cues used during the localization and navigation segments of the Motor Space Navigation (MSN) task.**

**(A)** Histogram showing number of cues most frequently sampled during the localization (LC) segment of the MSN task. Cues highlighted in yellow are the most frequently sampled.
**(B)** Heatmap indicating the frequency of presentation of cues in LC and during the navigation segment (C1/C2).
**(C)** Angular histogram showing the frequency of inferred trajectories from LC to C1/C2 in the MSN task.
**(D)** Performance of learners and non-learners in the MSN task across sessions. Error bars represent one std.

**SI - Additional Behavioral Assessments:**

To further evaluate participants' behavioral performance, we had participants perform a localization-only task at the end of Day 3, in which they executed an isometric exertion and directly selected the cue that matched the exerted force and time levels (**Supplementary Fig. 5 A, B**). Unlike the MSN task, the localization-only task directly tested the overt behavioral response following localization. Participants performed an isometric exertion that mirrored the localization segment of the MSN task, where the cue was replaced with an arbitrary circle. Resembling the MSN task, participants had to identify which of the 16 cues best matched the exertion. Following exertion, participants were presented with a screen that showed the correct cue as well as seven other distractor cues. Participants had to select the cue they thought best matched the exertion profile (See **Methods**). Participants who met the performance criteria in the MSN task were highly accurate in localizing cues (mean accuracy ± SE, 0.87 ± 0.02, **Supplementary Fig. 5C, D**), consistent with their ability to precisely link sensorimotor feedback with the learned force-time space.

To evaluate participants' internal representation of the force-time space, we administered a spatial reconstruction task on days 2 and 3. In this task, participants were presented with an empty 2D force-time map and had to reconstruct the full 4x4 force-time space (**Supplementary Fig. 6A, B**) (See **Methods**). Participants who met the performance criteria in the MSN task correctly reconstructed the full force-time space on both days (**Supplementary Fig. 6C**), indicative of a stable and coherent representation of the learned force-time space.

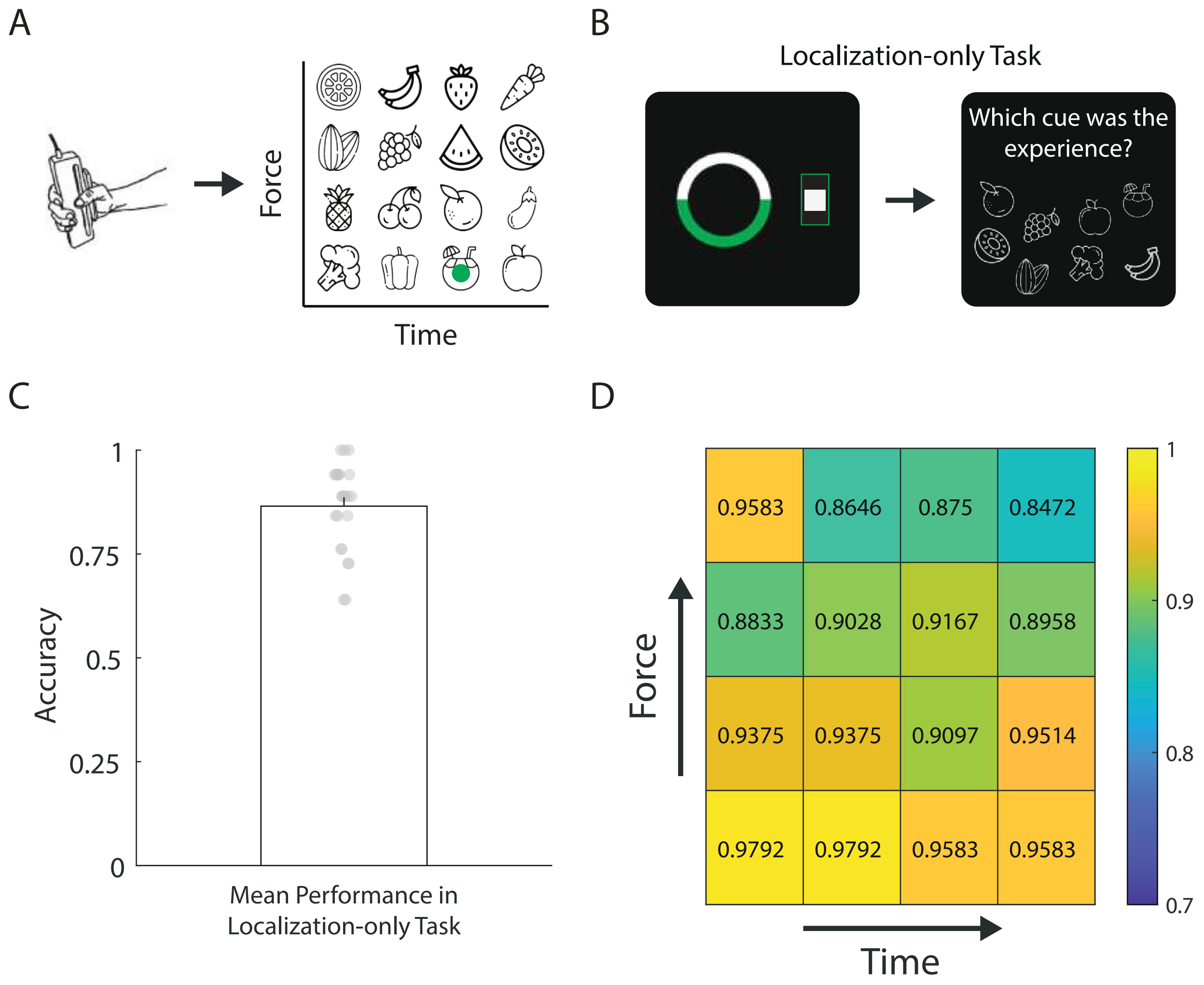


**Supplementary Figure 5: Performance of participants in the Localization-only task**

**(A)** Conceptual visualization of the localization-only task. Participants had to perform an exertion and identify the cue that best matched the force and time level of the exertion.
**(B)** In the localization-only task, participants first performed an exertion identifcal to the localization segment in the MSN task. They then were presented with a screen that showed the correct cue that matched the exertion profile in force and time level as well as seven other distractor cues. Participants had to select the correct cue. participants were provided feedback depending on whether they selected the correct or incorrect cue. If the incorrect cue was selected, that trial was sent to the end of the trial presentation. Participants had to correctly identify all 16 cues to finish the localization-only task.
**(C)** Mean performance in the localization-only task. Error bar represents one sem.
**(D)** Mean performance in the localization-only task for each of the 16 cues in the force-time space.

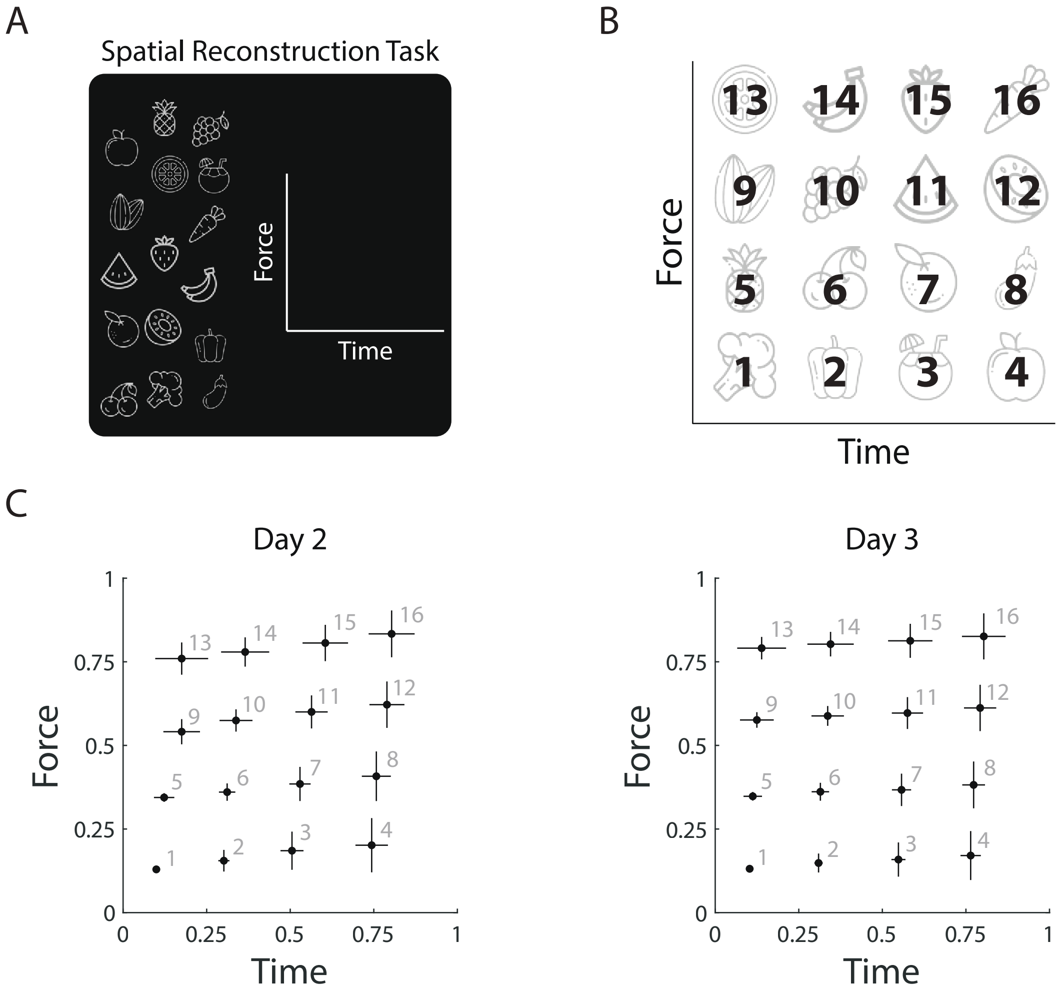


**Supplementary Figure 6: Performance of participants in the spatial reconstruction task.**

**(A)** In the spatial reconstruction task, participants were presented with the 16 cues that were organized in a randomized non-uniform way and an empty force-time space. Participants were instructed to complete the force-time space by dragging the cues to their respective location in the force-time space.
**(B)** For visualization purposes in subsequent plots, the cues were assigned labels from 1-16.
**(C)** Average placement location of each of the 16 cues in force-time space when participants performed the drag & drop task at the end of days 2 and 3. Error bars represent one sem.

$$\text{MSN Accuracy} \sim \text{Day} + \text{Phase} + \text{Reaction Time (RT)} + \Delta\ \text{Distance} (\Delta D) + \Sigma\ \text{Distance} (\Sigma D) + \text{Trajectory Similarity} (\theta) + \Delta D \text{ X Day} + \Sigma D \text{ X Day} + \theta \text{ X Day} + (1|\text{Participant})$$

| Fixed Effects Coefficients | Estimate | Standard Error | T | P Value |
|---|---|---|---|---|
| Intercept | 2.39 | 0.05 | 45.78 | < 0.001 |
| Day | 0.78 | 0.04 | 17.76 | < 0.001 |
| Phase | 0.31 | 0.03 | 9.52 | < 0.001 |
| Reaction Time (RT) | -0.32 | 0.03 | -10.78 | < 0.001 |
| Δ Distance (ΔD) | 0.84 | 0.05 | 17.30 | < 0.001 |
| Σ Distance (ΣD) | -0.34 | 0.04 | -8.04 | < 0.001 |
| Trajectory Similarity ($\theta$) | -0.34 | 0.05 | -6.80 | < 0.001 |
| ΔD X Day | 0.25 | 0.04 | 5.98 | < 0.001 |
| ΣD X Day | -0.20 | 0.04 | -5.30 | < 0.001 |
| $\theta$ X Day | -0.14 | 0.05 | -2.93 | 0.003 |

**Supplementary Table 1: Modeling trial-by-trial accuracy in the MSN task**

A mixed-effects logistic regression was used to model trial-by-trial accuracy (Correct = 1, Incorrect = 0) as a function of spatial metrics, experimental day (Day = 1,2,3,4), phase number (Before MSN training = 1, After MSN training = 2), reaction time, and the interaction between spatial metrics and experimental day. Participant ID was entered as random effects. All metrics were z-scored.

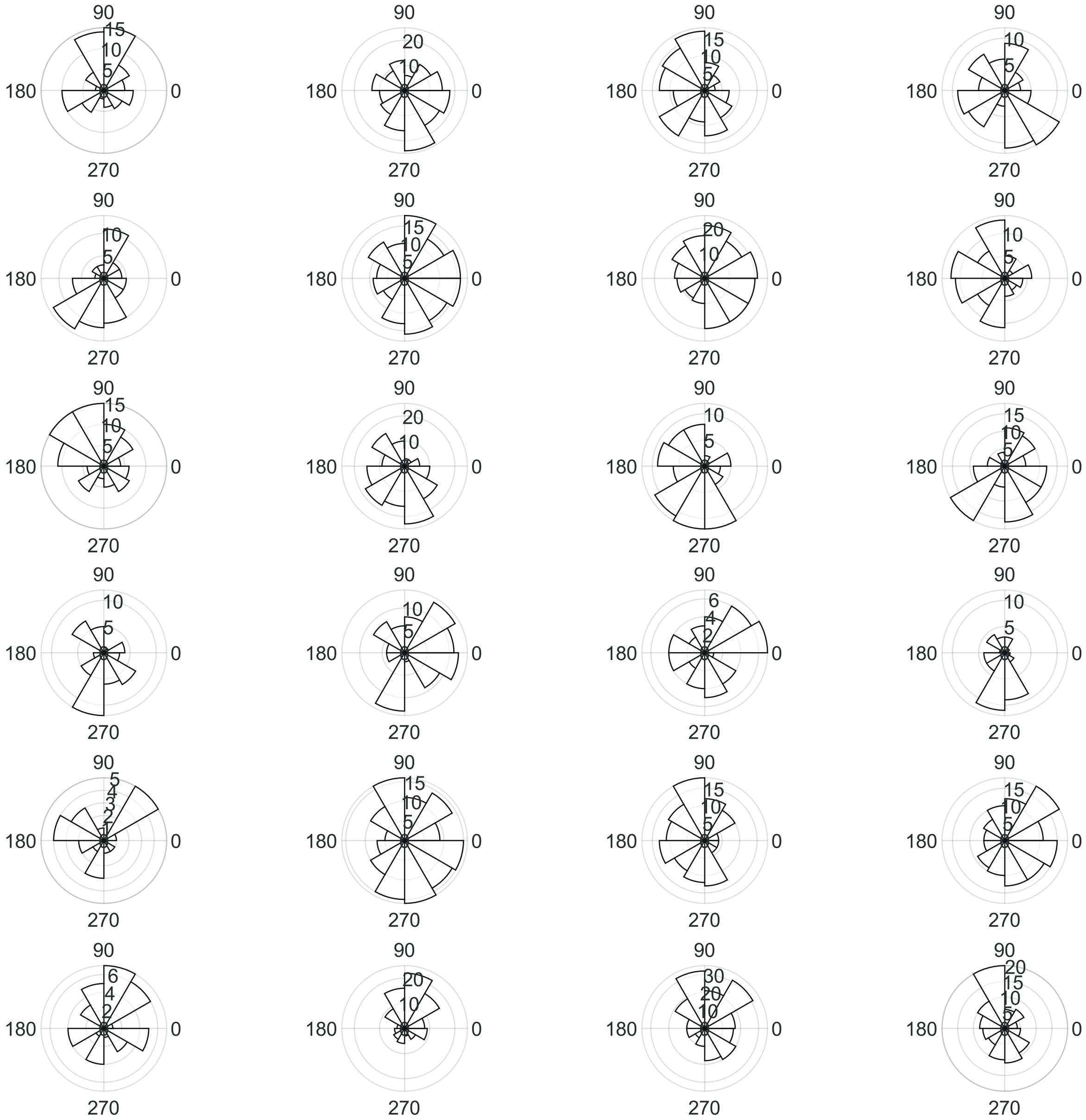

**Supplementary Figure 7: Nonuniform distribution of ERC grid orientation**

Polar histogram showing the estimated grid orientations for voxels within the ERC for each participant (n = 24). Grid orientations were significantly clustered in 18/24 participants (Rayleigh's test for nonunifromity; mean z ± SE = 5.37 ± 0.80).

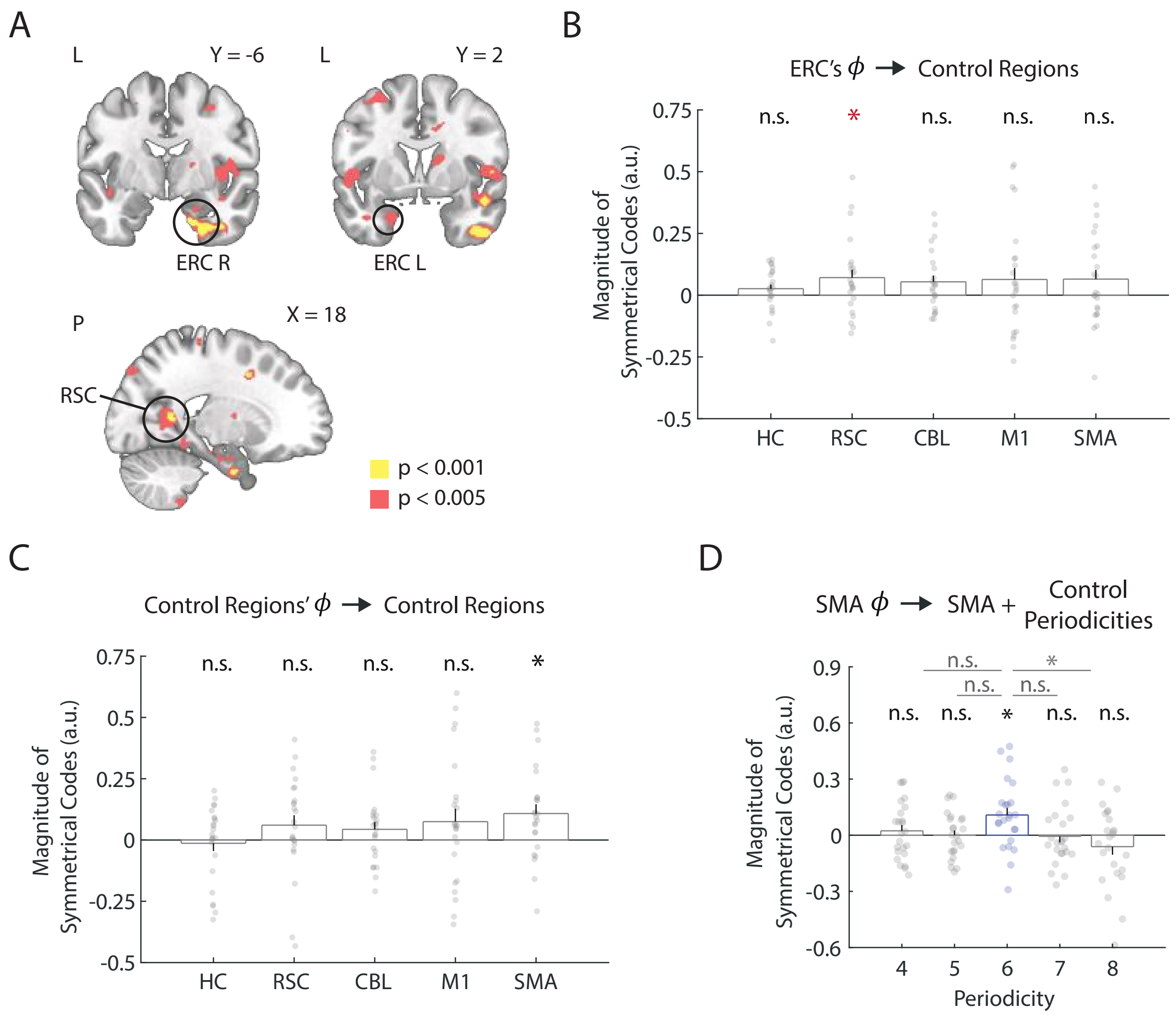


**Supplementary Figure 8: Testing for periodicity signals in control regions**

**(A)** Whole brain contrasts showing brain regions that were correlated with 6-fold periodicity signals using grid orientation estimated from the ERC. Contrast is displayed at $p < 0.001$ uncorrected (yellow) and $p < 0.005$ uncorrected (red).
**(B)** Activity in the control regions were tested for 6-fold periodicity signals using grid orientation estimated from the ERC. The RSC showed evidence of 6-fold modulation (* $p < 0.05$ uncorrected) which did not survive correction for multiple comparisons across number of tested ROIs (n = 5). Error bars represent one sem.
**(C)** Activity in the control regions were tested for 6-fold periodicity signals using grid orientation independently estimated from each region during the navigation phase. The supplementary motor area (SMA) displayed activity that was significantly correlated with 6-fold periodicity signals (* $p < 0.05$; two-tailed one sample Wilcoxon signed-rank test, corrected for number of regions (n = 5) using Holm-Bonferroni method.). The remaining four control regions did not display significant correlations with 6-fold periodicity signals (all $p > 0.05$). Error bars represent one sem.
**(D)** Activity in the supplementary motor area (SMA) was specifically correlated with 6-fold periodicity signals (* $p < 0.05$; two-tailed one sample Wilcoxon signed-rank test, corrected for number of periodicities (n = 5) using Holm-Bonferroni method.) but not with control periodicities (n = 4,5,7,8) ($p > 0.05$).

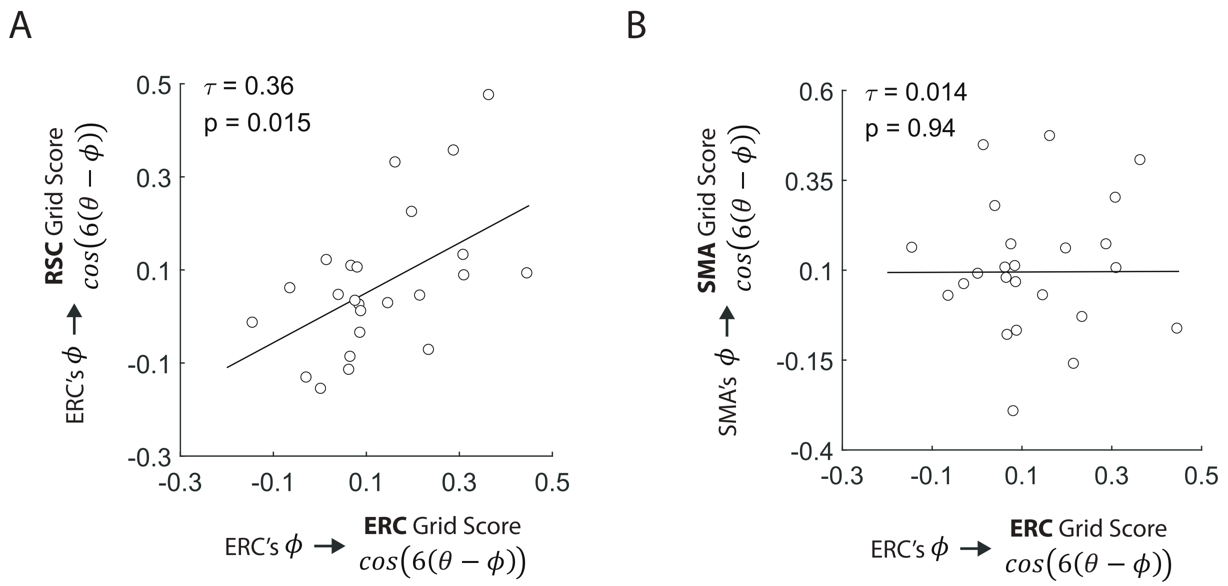


**Supplementary Figure 9: Quantifying relationship between grid-like signals from different brain regions**

**(A)** The magnitude of hexadirectional modulation in ERC and RSC (estimated using ERC's ϕ) were significantly correlated (Kendall's $\tau = 0.36$, $p = 0.015$), suggesting that individuals with stronger grid-like activity in ERC also showed greater grid-like modulation in RSC.
**(B)** The magnitude of hexadirectional modulation in ERC and SMA (estimated using SMA's ϕ) were not correlated (Kendall's $\tau = 0.014$, $p = 0.94$).

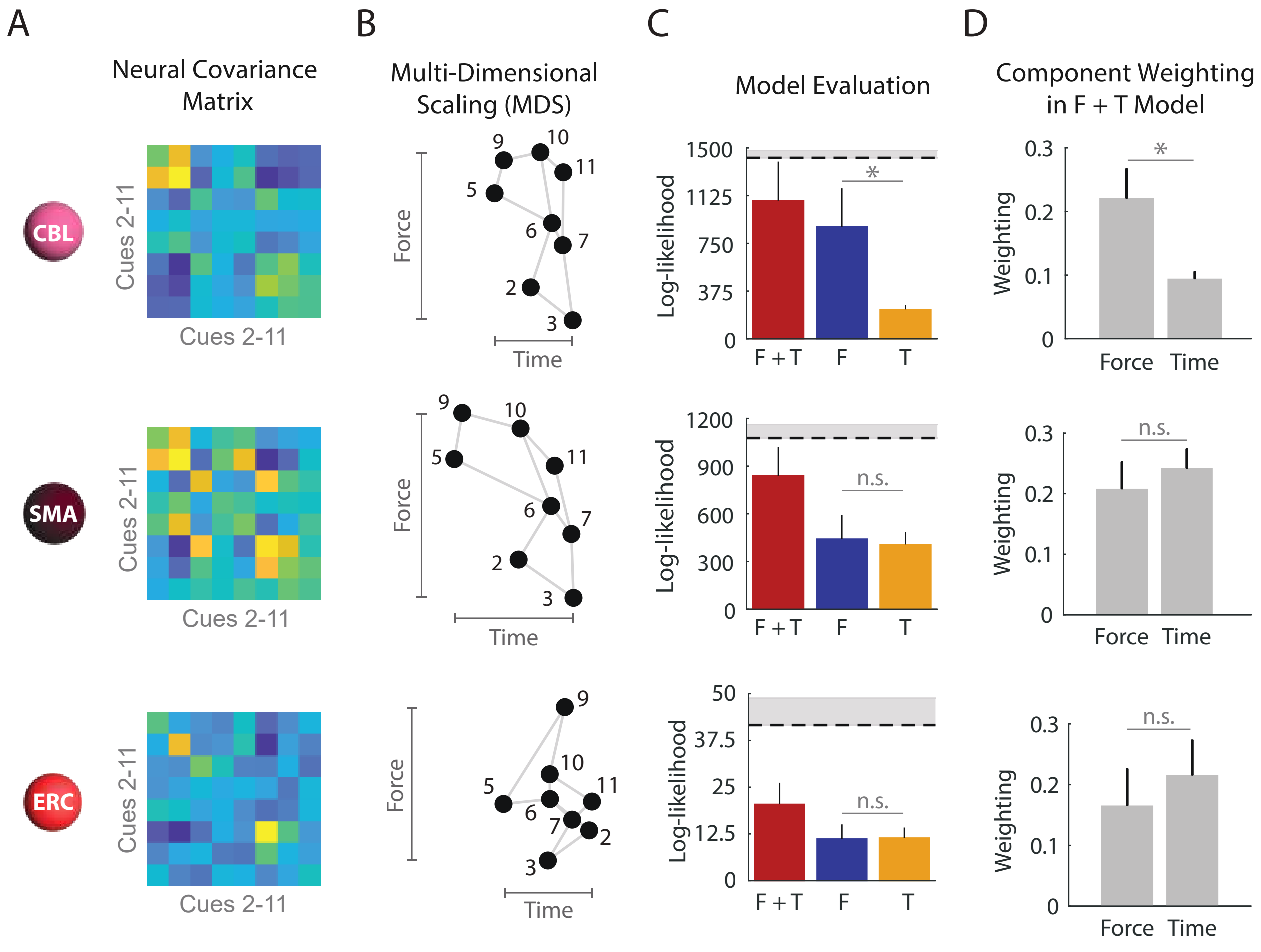


**Supplementary Figure 10: PCM results for the cerebellum, supplementary motor area, and entorhinal cortex**

**(A)** PCM results for the CBL (top), SMA (middle), and ERC (bottom). The cross-validated estimates of the neural second moment matrix describes the activity profile distribution for each brain region when exerting cues 2-11.
**(B)** Multi-Dimensional Scaling was performed (middle column) where the 8 cues were projected on the eigenvectors of the second moment matrix for each representative brain region.
**(C)** The performance of each model (right column) for predicting the neural second moment matrix was evaluated by computing model evidence based on the log-Bayes factor for the models against a null model. The force (F) and time (T) models predicted the neural second moment matrices using both F and T models. The force and time (F + T) model utilized both components. The dotted line represents the lower bound noise ceiling. * $p < 0.05$; two-tailed paired Wilcoxon signed-rank test, corrected for number of sensorimotor brain regions ($n = 3$) using Holm-Bonferroni method. Error bars represent one sem.
**(D)** The weights associated with the force and time components in the F + T model. * $p < 0.05$; two-tailed paired Wilcoxon signed-rank test, corrected for number of sensorimotor brain regions ($n = 3$) using Holm-Bonferroni method. Error bars represent one sem.

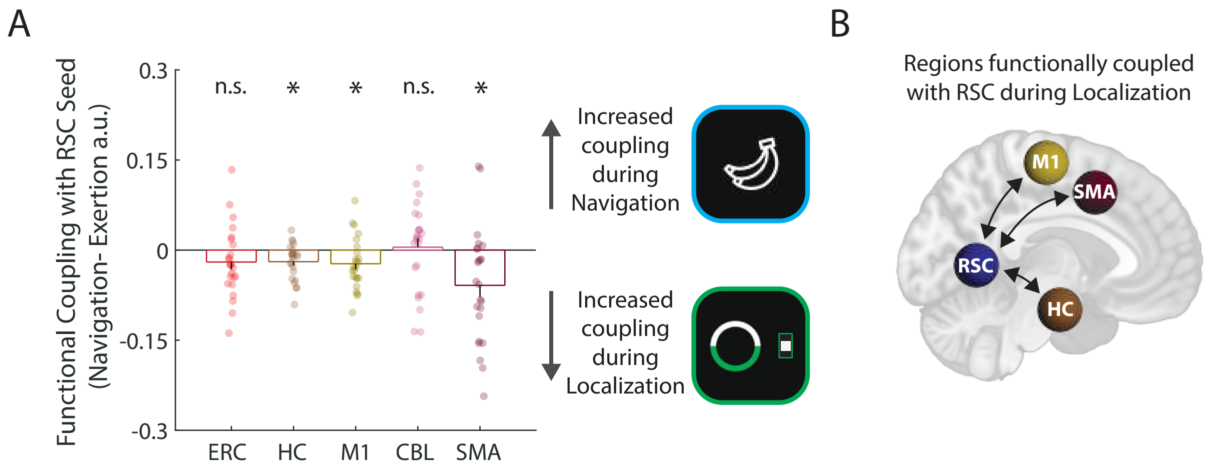


**Supplementary Figure 11: Psychophysiological interaction analysis results measuring changes in functional coupling with RSC at time of navigation vs. localization.**

**(A)** We performed a Psychophysiological interaction (PPI) analysis to determine which regions displayed significant functional coupling with the retrosplenial cortex (RSC) in the localization vs. navigation conditions. The RSC displayed significantly greater functional coupling with hippocampus (HC), primary motor cortex (M1), and supplementary motor area (SMA) in the localization condition. Error bars represent one sem. * $P < 0.05$; two-tailed one sample Wilcoxon signed-rank test, corrected for number of regions ($n = 5$) using Holm-Bonferroni method.

**(B)** Visualization of PPI results which shows functional coupling of RSC with sensorimotor (M1, SMA) and mnemonic (HC) region(s).

**SI - Dynamic Causal Modeling:**

Building on the observed functional coupling, we used dynamic causal modeling (DCM) to examine directed causal interactions between the sensorimotor and mnemonic brain regions. We specified a deterministic, bilinear DCM to test directed influences between the RSC, M1, SMA, and HC (i.e., effective connectivity), which we chose as seed regions based on our PPI findings. We also assessed how connectivity measures changed during the localization and navigation conditions (**Supplementary Fig. 12A**). Based on our initial GLM results (**Supplementary Fig. 7**), we allowed the navigation condition to directly influence activity in mnemonic regions (HC, RSC) and the localization condition to directly influence activity in sensorimotor regions (M1, SMA) (**Supplementary Fig. 12A**). We employed the Parametric Empirical Bayes (PEB) framework, estimating a single 'full' model that included all relevant connections for each participant (1st level), then examining commonalities and differences in connection parameters across participants (2nd level). Importantly, the sign of each connection indicates whether a region's activity is boosted (positive/excitatory) or suppressed (negative/inhibitory) by the experimental factors or inputs from other regions. Following PEB guidelines, we performed an iterative search over reduced models—where specific connections were deactivated—to evaluate each connection's influence on model fit using their posterior probability (Pp) (**Supplementary Fig. 12B**).

Given the large number of connection parameters, we interpreted connections with Pp greater than 0.99, which indicated very strong evidence (**Supplementary Fig. 12B, C**). During localization, we identified extensive modulatory connections between mnemonic and sensorimotor regions (**Supplementary Fig. 12C; Supplementary Table 2**). Consistent with our PPI findings, the RSC received inputs from M1, SMA, and HC. DCM revealed additional interactions not detected by the PPI results, including modulatory connections between the HC and sensorimotor regions. Notably, the nature of these connections varied by region type: connections originating from sensorimotor regions were consistently inhibitory, while those from mnemonic regions were excitatory (**Supplementary Fig. 12C**). During navigation, however, only the connection from HC to SMA showed very strong evidence (Pp > 0.99), suggesting a general functional decoupling between mnemonic and sensorimotor regions once the cue-exertion association had already been established in the navigation segment (**Supplementary Fig. 12D; Supplementary Table 3**).

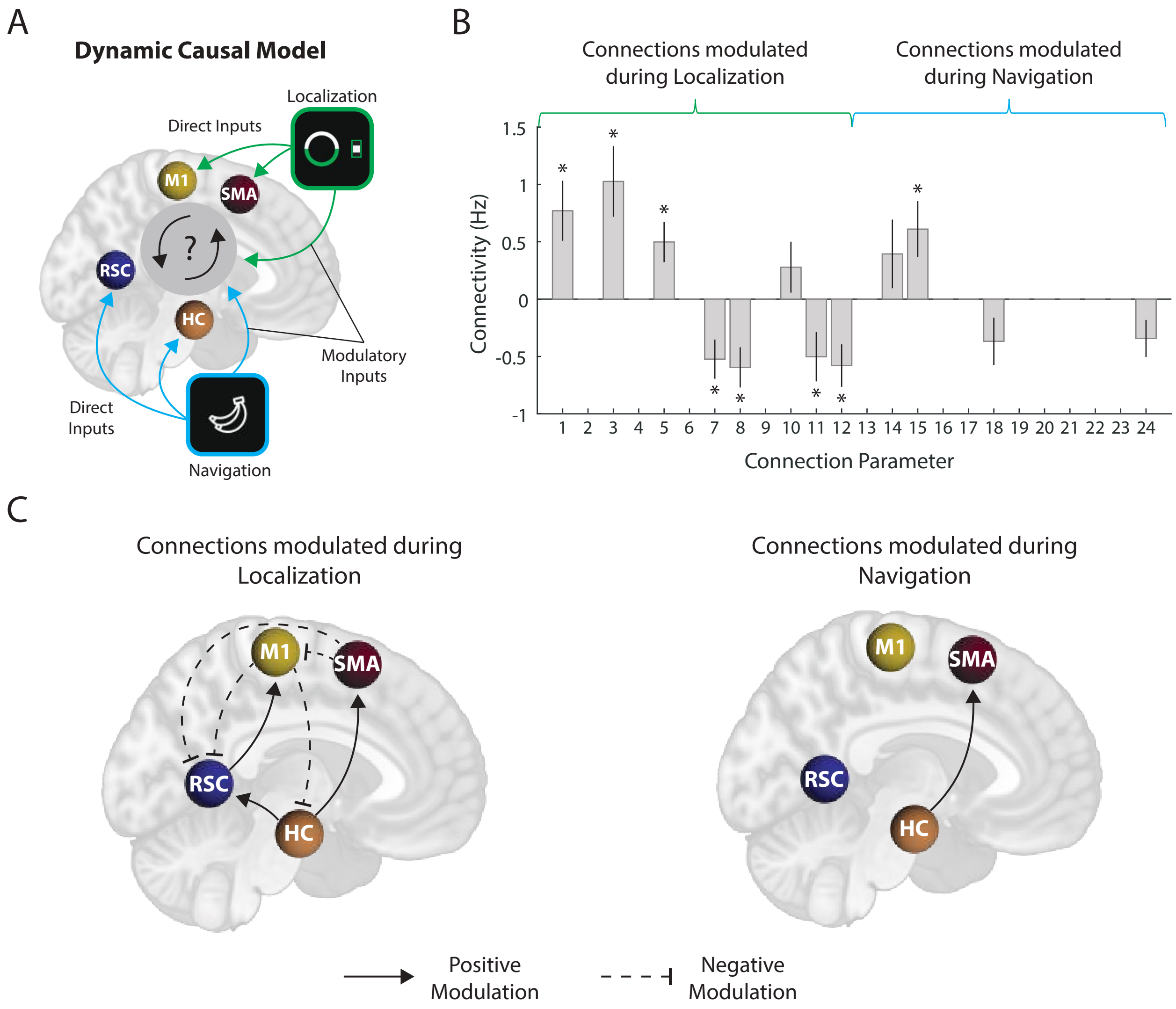


**Supplementary Figure 12: Dynamic causal modeling shows changes in effective connectivity between sensorimotor and mnemonic brain regions at time of localization and navigation**

**(A)** We created a dynamic causal model (DCM) where we assigned RSC, HC, M1 and SMA as a priori seed regions of interest which was informed by what we find from the PPI analysis (Supplementary Fig. 11). We enabled all bi-directional connections between each region. Based on the preliminary GLM results (Supplementary Fig. 7), we enabled the localization condition to directly drive activity in M1 and SMA and for the navigation condition to directly drive activity in RSC and HC (Direct inputs). We then enabled both localization and navigation conditions to modulate all bi-directional connections between each region (Modulatory Inputs).
**(B)** We applied the parametric empirical bayes (PEB) framework where following the estimation of a full DCM model that enables all possible connections between regions, Bayesian model reduction (BMR) was used to compare evidence for reduced models and iteratively discard parameters that do not contribute to model evidence. This process stopped when discarding any parameter began to decrease model evidence. Connection parameters that survive this iterative search along with their connectivity estimate is shown in the plot. Of the 256 models that survived the iterative search, the posterior probability of each connection parameter was computed by performing a Bayesian model comparison to compute the difference in model evidence of the PEB model with the parameter turned on versus the equivalent PEB model with the parameter turned off. We interpreted parameters with posterior probability greater than 0.99 (*), which corresponded to very strong evidence for connections modulated during localization (1-12) and navigation (13-24). Error bars represent 90% credible interval.
**(C)** Visualization of DCM parameters that show very strong evidence of connections being modulated during localization (left) and navigation (right). Solid arrows indicate positive modulation whereas dashed flat arrows indicate negative modulation.

| Connection Parameter | Source | Target | Connectivity (Hz) | Posterior Probability |
|---|---|---|---|---|
| 1 | HC | RSC | 0.77 | 1.00 |
| 3 | HC | SMA | 1.03 | 1.00 |
| 5 | RSC | M1 | 0.50 | 1.00 |
| 7 | M1 | HC | -0.52 | 1.00 |
| 8 | M1 | RSC | -0.60 | 1.0 |
| 10 | SMA | HC | 0.28 | 0.90 |
| 11 | SMA | RSC | -0.50 | 1.00 |
| 12 | SMA | M1 | -0.58 | 1.00 |

**Supplementary Table 2: DCM connection parameters modulated during the Localization condition.**

| Connection Parameter | Source | Target | Connectivity (Hz) | Posterior Probability |
|---|---|---|---|---|
| 14 | HC | M1 | 0.39 | 0.92 |
| 15 | HC | SMA | 0.61 | 1.00 |
| 18 | RSC | SMA | -0.368 | 0.98 |
| 24 | SMA | M1 | -0.343 | 0.99 |

**Supplementary Table 3: DCM connection parameters modulated during the Navigation condition.**

$$\text{MSN Accuracy} \sim 1 - 0.5e^{-bs} \quad b = \text{Learning rate}, \quad s = \text{Session}$$

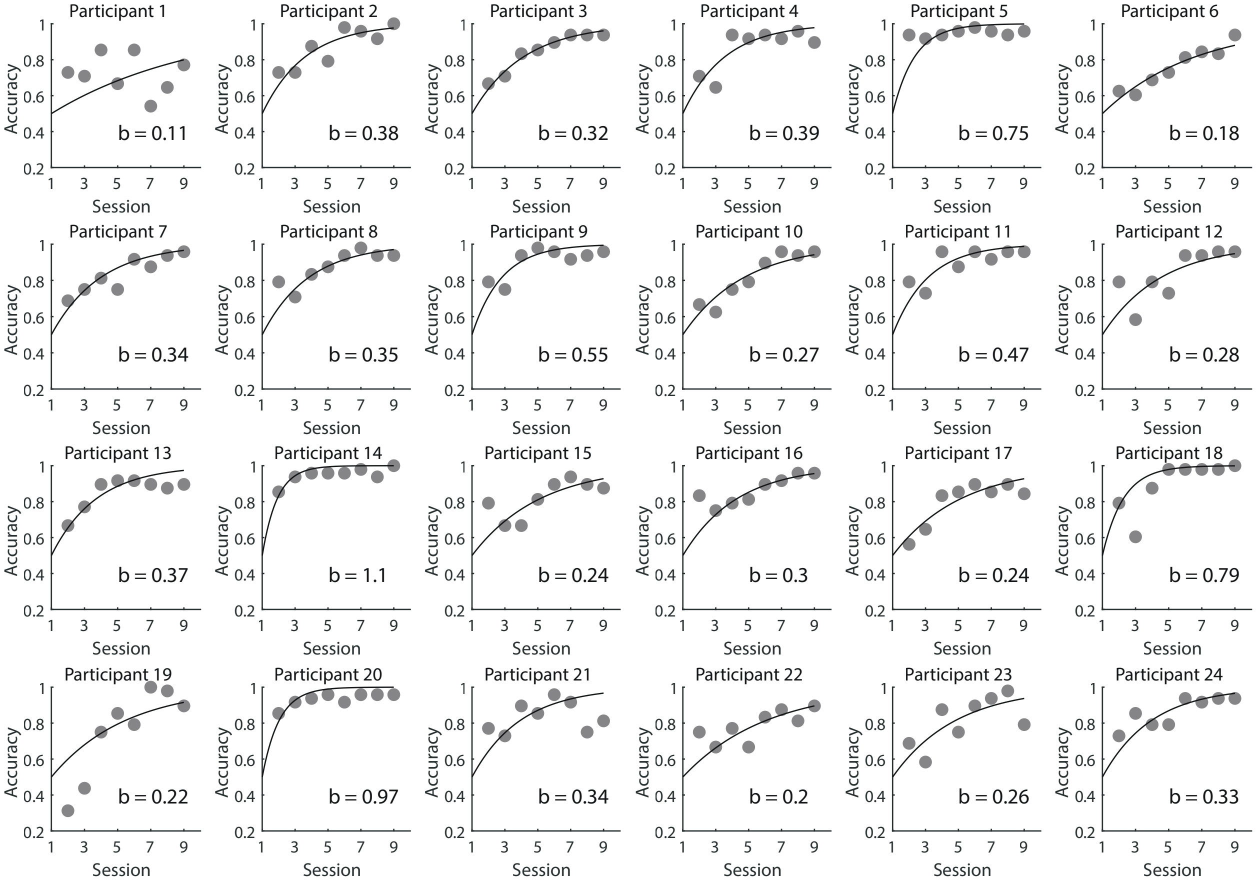


**Supplementary Figure 13: Individual fitting of participants' performance in the MSN task across sessions**

Individual fitting of each participants' improvement in the MSN task using an exponential function. Participants who improved and stabilized faster in the MSN task had a greater b parameter than those who learned more gradually (lower b parameter).

A

Mean Warping Mnemonic Regions (RSC, HC, ERC) ~ Learning Rate (b) + Perceptual Bias (ΔForce - ΔTime)

| Fixed Effects Coefficients | Estimate | Standard Error | T | P Value |
|---|---|---|---|---|
| Intercept | 0.54 | 0.04 | 14.23 | < 0.001 |
| Learning Rate (b) | -0.13 | 0.04 | -3.29 | 0.0034 |
| Perceptual Bias (ΔForce -ΔTime) | 0.12 | 0.04 | 3.10 | 0.0054 |

B

Mean Warping Sensorimotor Regions (M1, CBL, SMA) ~ Learning Rate (b) + Perceptual Bias (ΔForce - ΔTime)

| Fixed Effects Coefficients | Estimate | Standard Error | T | P Value |
|---|---|---|---|---|
| Intercept | 0.47 | 0.05 | 10.34 | < 0.001 |
| Learning Rate (b) | -0.03 | 0.05 | -0.56 | 0.58 |
| Perceptual Bias (ΔForce -ΔTime) | 0 | 0.05 | -0.06 | 0.95 |

C

Warping in RSC ~ Learning Rate (b) + Perceptual Bias (ΔForce - ΔTime)

| Fixed Effects Coefficients | Estimate | Standard Error | T | P Value |
|---|---|---|---|---|
| Intercept | 0.56 | 0.06 | 9.63 | < 0.001 |
| Learning Rate (b) | -0.16 | 0.06 | -2.66 | 0.01 |
| Perceptual Bias (ΔForce -ΔTime) | 0.13 | 0.06 | 2.22 | 0.04 |

D

Warping in HC ~ Learning Rate (b) + Perceptual Bias (ΔForce - ΔTime)

| Fixed Effects Coefficients | Estimate | Standard Error | T | P Value |
|---|---|---|---|---|
| Intercept | 0.48 | 0.07 | 7.42 | < 0.001 |
| Learning Rate (b) | -0.16 | 0.07 | -2.43 | 0.02 |
| Perceptual Bias (ΔForce -ΔTime) | 0.05 | 0.07 | 0.77 | 0.45 |

E

Warping in ERC ~ Learning Rate (b) + Perceptual Bias (ΔForce - ΔTime)

| Fixed Effects Coefficients | Estimate | Standard Error | T | P Value |
|---|---|---|---|---|
| Intercept | 0.59 | 0.07 | 8.19 | < 0.001 |
| Learning Rate (b) | -0.07 | 0.07 | -0.92 | 0.37 |
| Perceptual Bias (ΔForce -ΔTime) | 0.18 | 0.07 | 2.40 | 0.03 |

**Supplementary Figure 14: Robust linear regression results for predicting warping in mnemonic and sensorimotor regions using learning rate and perceptual bias**

**(A)** Robust linear regression was used to model each participants' mean warping in mnemonic regions (RSC, HC, ERC) as a function of their learning rate (b) and perceptual bias (ΔForce - ΔTime).
**(B)** The same process in (A) was used to model mean warping in sensorimotor regions (M1, CBL, SMA)
**(C)** The same process in (A) was used to model mean warping in just the RSC.
**(D)** The same process in (A) was used to model mean warping in just the HC.
**(E)** The same process in (A) was used to model mean warping in just the ERC.

A

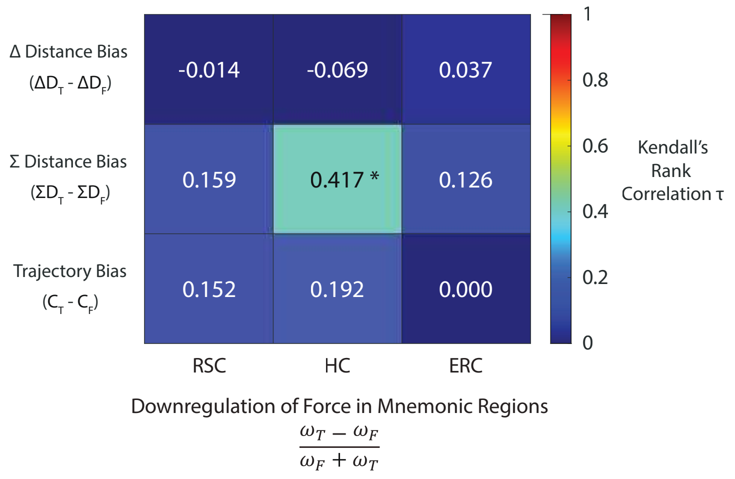


B

Σ Distance Bias ($\Sigma D_T - \Sigma D_F$) ~ RSC Force Downregulation + HC Force Downregulation + ERC Force Downregulation

| Fixed Effects Coefficients | Estimate | Standard Error | T | P Value |
|---|---|---|---|---|
| Intercept | 0.01 | 0.06 | 0.15 | 0.88 |
| RSC $\frac{\omega_T - \omega_F}{\omega_F + \omega_T}$ | -0.10 | 0.11 | -0.95 | 0.36 |
| HC $\frac{\omega_T - \omega_F}{\omega_F + \omega_T}$ | 0.30 | 0.11 | -2.81 | 0.01 |
| ERC $\frac{\omega_T - \omega_F}{\omega_F + \omega_T}$ | 0.05 | 0.08 | 0.65 | 0.52 |

C

Accuracy in Localization-Only Task ~ RSC Warping + HC Warping + ERC Warping

| Fixed Effects Coefficients | Estimate | Standard Error | T | P Value |
|---|---|---|---|---|
| Intercept | 0.97 | 0.05 | 21.46 | < 0.001 |
| RSC $\left\lvert\frac{\omega_T - \omega_F}{\omega_F + \omega_T}\right\rvert$ | -0.22 | 0.08 | -2.77 | 0.01 |
| HC $\left\lvert\frac{\omega_T - \omega_F}{\omega_F + \omega_T}\right\rvert$ | 0.11 | 0.07 | 1.45 | 0.16 |
| ERC $\left\lvert\frac{\omega_T - \omega_F}{\omega_F + \omega_T}\right\rvert$ | -0.03 | 0.05 | -0.48 | 0.64 |

D

RSC Force Downregulation ~ Force Error + Time Error

| Fixed Effects Coefficients | Estimate | Standard Error | T | P Value |
|---|---|---|---|---|
| Intercept | 0.06 | 0.16 | 0.37 | 0.72 |
| Force Error | 2.76 | 1.17 | 2.36 | 0.03 |
| Time Error | -0.52 | 1.61 | -0.33 | 0.75 |

**Supplementary Figure 15: The HC and RSC reflect distinct computational processes when navigating sensorimotor space**

**(A)** Kendall's rank correlation matrix detailing the correlation between spatial bias metrics (y axis) and the downregulation of force in mnemonic regions (x-axis). Kendall's tau ($\tau$) is displayed for each correlation. * $p < 0.05$, family-wise error (FWE) corrected over number of correlations ($n = 9$)
**(B)** $\Sigma$ Distance Bias was modeled using robust linear regression as a function of force downregulation in RSC, HC and ERC for each participant.
**(C)** Participants' accuracy in the localization-only task was modeled using robust linear regression as a function of warping in RSC, HC and ERC.
**(D)** Force downregulation in RSC was modeled using robust linear regression as a function of force and time error.
